# STING Licensing of Type I Dendritic Cells Potentiates Antitumor Immunity

**DOI:** 10.1101/2024.01.02.573934

**Authors:** Jian Wang, Suxin Li, Maggie Wang, Xu Wang, Shuqing Chen, Zhichen Sun, Xiubao Ren, Gang Huang, Baran D. Sumer, Nan Yan, Yang-Xin Fu, Jinming Gao

## Abstract

Stimulator of interferon genes (STING) is an immune adaptor protein that senses cyclic GMP-AMP (cGAMP) in response to self or microbial cytosolic DNA as a danger signal. STING is ubiquitously expressed in diverse cell populations including cancer cells with distinct cellular functions such as activation of type I interferons, autophagy induction, or triggering apoptosis. It is not well understood whether and which subsets of immune cells, stromal cells, or cancer cells are particularly important for STING-mediated antitumor immunity. Here using a polymeric STING-activating nanoparticle (PolySTING) with a “shock-and-lock” dual activation mechanism, we show type 1 conventional dendritic cell (cDC1) is essential for STING-mediated rejection of multiple established and metastatic murine tumors. STING status in the host but not in the cancer cells (*Tmem173^-/-^*) is important for antitumor efficacy. Specific depletion of cDC1 (*Batf3^-/-^*) or STING deficiency in cDC1 (*XCR1^cre^STING^fl/fl^*) abolished PolySTING efficacy, whereas depletion of other myeloid cells had little effect. Adoptive transfer of wildtype cDC1 in *Batf3^-/-^* mice restored antitumor efficacy while transfer of cDC1 with STING or IRF3 deficiency failed to rescue. PolySTING induced a specific chemokine signature in wildtype but not *Batf3^-/-^* mice. Multiplexed immunohistochemistry analysis of STING-activating cDC1s in resected tumors correlates with patient survival while also showing increased expressions after neoadjuvant pembrolizumab therapy in non-small cell lung cancer patients. Therefore, we have defined that a subset of myeloid cells is essential for STING-mediated antitumor immunity with associated biomarkers for prognosis.

**One Sentence Summary:** A “shock-and-lock” nanoparticle agonist induces direct STING signaling in type 1 conventional dendritic cells to drive antitumor immunity with defined biomarkers

## INTRODUCTION

STING (aka MITA, MYPS and encoded by *TMEM173* gene) is an endoplasmic reticulum (ER)-associated signaling protein that is essential for transcriptional regulation of numerous host defense genes against infection and cancer (*1-3*). STING is activated by 2’, 3’-cyclic guanosine monophosphate-adenosine monophosphate (cGAMP) (*4, 5*), an endogenous secondary messenger, which is produced by cGAMP synthase (cGAS) in response to microbial or self-DNA as a danger signal (*6*). Upon cGAMP binding, STING homodimer undergoes extensive conformational change that leads to oligomerization on the ER surface and translocation to the Golgi. STING oligomerization recruits and activates TANK-binding kinase 1 (TBK1) and inhibitor of κB kinases (IKKs), which in turn activate interferon regulatory factor 3 (IRF3) and nuclear factor κB (NFκB), respectively. IRF3 and NFκB cooperate to induce type I interferons and other inflammatory cytokines to mount a robust innate immune response. Besides immune activation, STING also triggers autophagy as a primordial function independent of TBK1 and IKK (*7*). Furthermore, high magnitude of STING signaling drives cell apoptosis in hepatocytes and lymphocytes through elevated ER stress (*8, 9*). Understanding the cell context of STING activation and resulting molecular responses is paramount to mount a protective immune response against cancer.

In this study, we leveraged a polymeric STING-activating nanoparticle (PolySTING, Fig. 1A) (*10*) to investigate the role of immune cell subset in STING-mediated antitumor immunity. PolySTING combines a burst (shock) STING activation by cGAMP followed by a sustained (lock) effect from a synthetic polymer, PSC7A. PolySTING displays tropism toward myeloid cells with minimal uptake in T cells to avoid STING-mediated T cell death. The cell tropism and “shock-and-lock” STING activation by PolySTING allow the identification of conventional type 1 dendritic cells (cDC1s) as a key cell driver for the rejection of malignant tumors. Our findings reveal direct targeting of cDC1s by a dual activating STING agonist offers a promising approach for STING-mediated cancer immunotherapy.

**Fig. 1.**
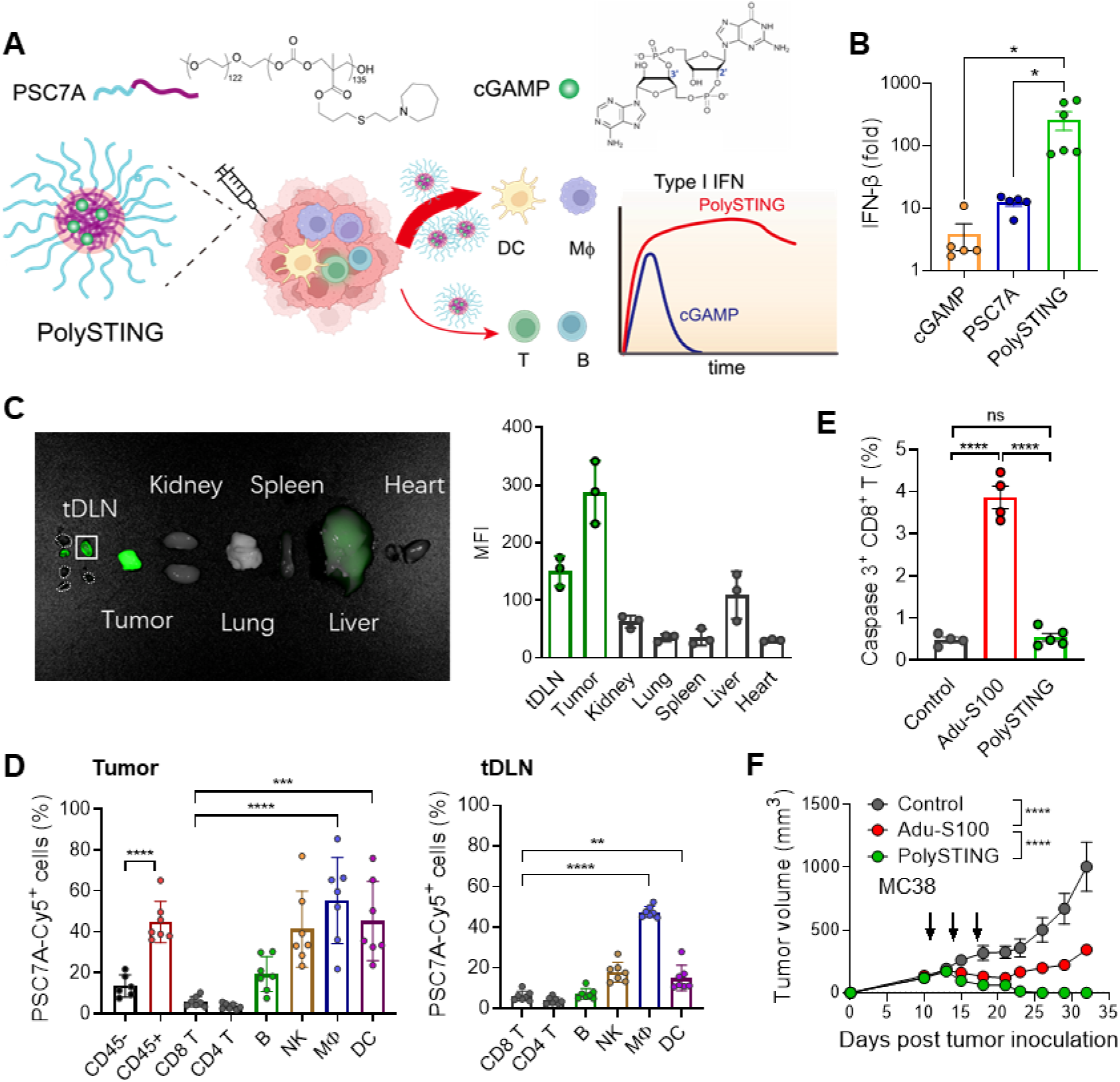
Design and characterization of a polymeric STING-activating nanoparticle (PolySTING) for myeloid tropic STING activation. (**A**) Schematic overview of PolySTING micelle nanoparticles with dual STING activation by 2’, 3’-cGAMP and PSC7A polymer. cGAMP is encapsulated in micelles self-assembled from pH-sensitive PSC7A polymers. (**B**) IFN-β-luciferase expressions in ISG-THP1 cells after free cGAMP, PSC7A polymer, or PolySTING treatment over 24 hours. Equivalent cGAMP concentrations (5 μM) were used in these studies. (**C**) *Ex vivo* imaging and tissue quantification of fluorescence intensity in MC38 tumors 24 hours post intratumoral administration of Cy5-labeled PolySTING. (**D**) Percentage of Cy5^+^ cells amongst cell populations in the MC38 tumors and tumor-draining lymph nodes following intratumoral administration of Cy5-labeled PolySTING. (**E**) Percentage of caspase 3^+^ CD8^+^ T cells after intratumoral administration of PolySTING or Adu-S100 at the same dose (50 μg per mouse). (**F**) Antitumor efficacy of PolySTING or Adu-S100 in MC38 tumor bearing mice. All the data in are represented as mean±sem, n≥5 in B,D,E,F and n=3 in C. Statical significance was calculated by two-tail t-test in B,D,E, or two-way ANOVA in F. ns: no significant difference, *: p<0.05, **: p<0.01, ***: p<0.001, ****: p<0.0001.

## RESULTS

### PolySTING is a potent STING agonist against hot and cold tumors

We produced PolySTING nanoparticles by encapsulating cGAMP in a micelle nanoparticle self-assembled from a STING-activating polymer, polyethylene oxide-*b*-PSC7A (PSC7A, Fig. 1A and fig. S1A). Previous work shows canonical STING activation by cGAMP alone led to rapid STING activation in 6 hours followed by degradation in lysosomes and loss of interferon expression after 12 hours, whereas polymer-induced STING condensation and activation prolonged the expression of type I interferons over 48 hours (*10*). After cGAMP loading in PSC7A micelles, dynamic light scattering analysis shows a diameter of 25±3 nm, and transmission electron microscopy displays a spherical morphology (fig. S1A). After incubation in ISG-THP1 cells over 24 hours, PolySTING produced robust IFN-β expression (>250-fold over saline control) compared to cGAMP (3-fold) or PSC7A NP (12-fold) groups (Fig. 1B). Time-dependent cytokine analysis illustrates the “shock-and-lock” STING activation profile from PolySTING compared to a transient response by cGAMP (fig. S1B).

We used Cy5-labeled PolySTING to quantify the nanoparticle biodistribution after intratumoral administration in MC38 colon carcinoma. Abundant accumulation of PolySTING was found in the tumor and tumor-draining lymph nodes (tDLN) at 24 hours over other organs (Fig. 1C). Flow cytometry analysis shows significantly higher cell uptake in the immune cells (CD45^+^) over cancer cells (CD45^-^) in MC38 tumors (Fig. 1D). Further examination of subsets of immune cell populations displayed significant tropism of PolySTING toward the dendritic cells, macrophages, and natural killer cells over T cells in the tumor microenvironment and tDLNs. In contrast, FITC-labelled cGAMP did not show dramatic cell tropism across immune cell populations in tumor tissues, and only negligible cGAMP signal (<1%) was found in tDLNs (fig. S1C). *In vitro* studies show bone marrow-derived dendritic cells (BMDCs) or macrophages (BMDMs) readily endocytose PolySTING with increased cGAMP uptake over free cGAMP (fig. S1, D to F). Intratumoral injection of small molecule STING agonists (e.g., Adu-S100) have been reported to cause T cell apoptosis and loss of T cell memory at high doses (*11*). For this purpose, we evaluated the effect of PolySTING (50 μg) versus Adu-S100 (50 μg) on T cell viability and tumor growth inhibition. Data show significantly increased apoptosis in CD8^+^ T cells 24 hours after one injection of Adu-S100 compared to PolySTING (Fig. 1E and fig. S1G). Furthermore, MC38 tumors regress after initial treatment by Adu-S100 followed by relapse, whereas PolySTING treatment led to complete tumor eradication (Fig. 1F). Additional dose-response studies show PolySTING achieved a much broader therapeutic window from 20 to 100 μg for tumor eradication, whereas Adu-S100 had the best antitumor effect at 20 μg (2/5 tumor free) (fig. S1, H and I).

PolySTING displays broad antitumor efficacy with prolonged survival over free cGAMP or glucose controls in immunogenic (e.g., MC38, human papilloma virus E6/E7-transfected TC-1 tumor, Fig. 2, A and B, and fig. S2, A and B) and immune-cold murine tumors (e.g., B16F10 melanoma, Lewis lung LLC2 carcinoma, Fig. 2, C and D, and fig. S2, C and D). To further explore if local treatment can generate sufficient T cells against metastasis, we selected triple negative 4T1 orthotopic mammary carcinoma that will have spontaneous lung metastasis starting on second week after inoculation. PolySTING monotherapy on day 11 not only inhibited the primary tumor growth, but also generated systemic immunity against the formation of metastatic 4T1 foci in the lung (Fig. 2E and fig. S2E). Impressively, PolySTING treatment is more potent than anti-PD-1 treatment for reducing metastasis. In the MC38 and B16F10 distal tumor models, PolySTING also drives growth inhibition in distal tumors after injection in the primary tumors (Fig. 2F, and fig. S3, A and B). Furthermore, combination of PolySTING with anti-PD-1 exhibited therapeutic synergy in MC38 distal tumors. Mice cured by PolySTING displayed long-term T cell memory as shown by the MC38 rechallenging experiment at both the healed primary tumor location and the contralateral site (Fig. 2G and fig. S3C). Therefore, local treatment of PolySTING generates potent systemic immunity at distal and metastatic sites with long-term memory effect.

**Fig. 2.**
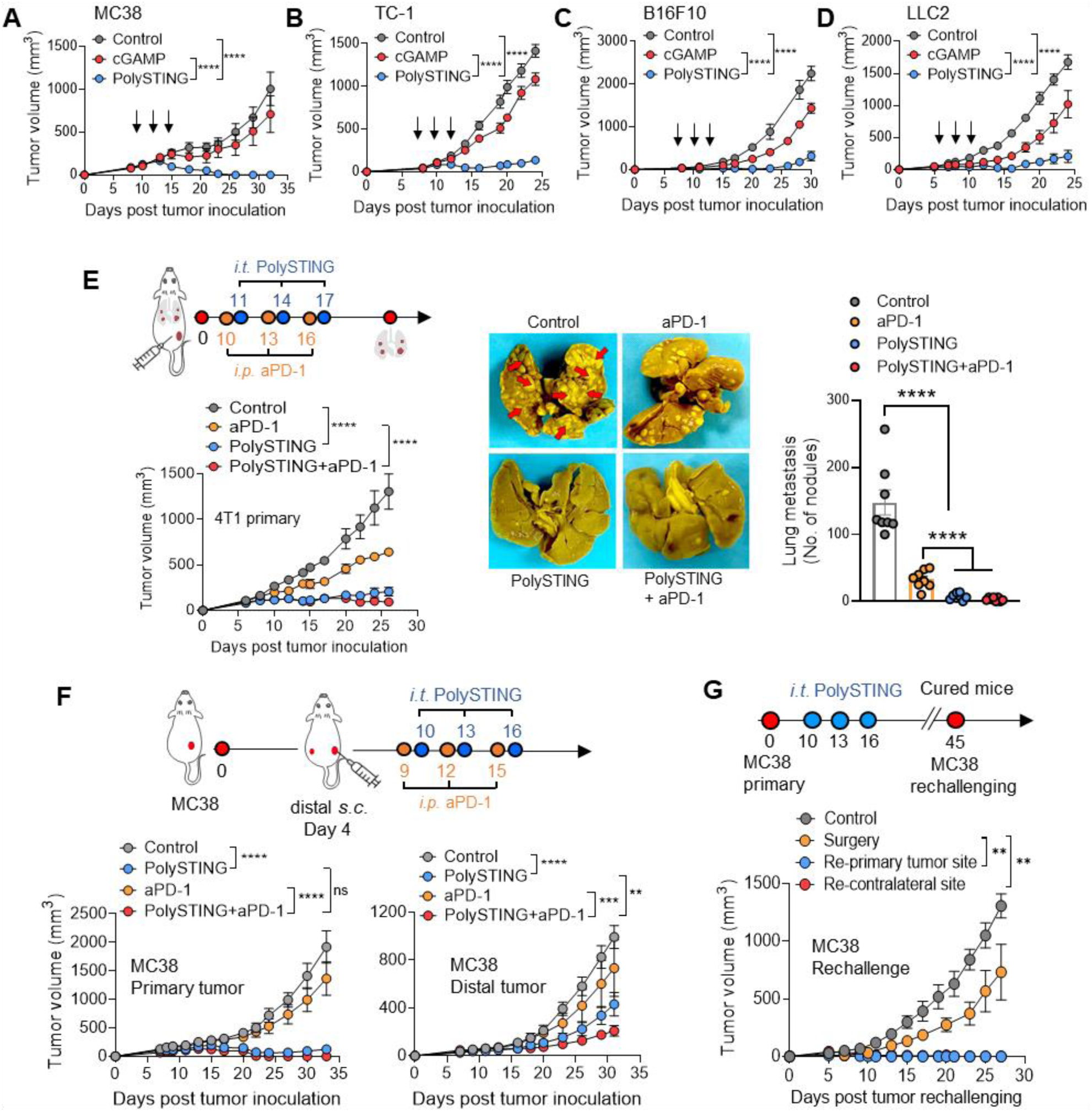
PolySTING achieves broad antitumor efficacy against established, metastatic, and distal tumors. **(A-D)** Intratumoral administration of PolySTING led to stronger growth inhibition of MC38, TC-1, B16F10, or LLC2 tumors over free cGAMP in C57BL/6 mice. (**E**) Primary tumor growth and metastasis to the lung in Balb/c mice bearing orthotopic triple negative 4T1 breast tumor after PolySTING and anti-PD-1 treatment. (**F**) Injected primary and untreated contralateral tumor growth in C57BL/6 mice bearing MC38 tumors after PolySTING and anti-PD-1 treatment. (**G**) MC38 tumor-bearing mice cured by PolySTING or surgical resection were re-challenged with MC38 on the primary tumor site or contralateral flank 45 days after inoculation. All the data are represented as mean±sem, n≥5. Statical significance was calculated by two-way ANOVA, ns: no significant difference, **: p<0.01, ***: p<0.001, ****: p<0.0001.

### Host STING-IFN-I signaling is important for PolySTING-mediated antitumor efficacy

We investigated the role of STING in the host versus MC38 cancer cells in antitumor immunity. We compared the tumor growth inhibition using STING wildtype MC38 cancer cells in host *Tmem173^−/−^* (STING knockout) mice, and *Tmem173^−/−^* MC38 cells in STING wildtype mice. PolySTING-mediated tumor eradiation was abolished in host *Tmem173^-/-^*animals with MC38 tumors but was retained in *Tmem173^−/−^* MC38 tumors with wildtype mice (Fig. 3, A to D). The predominant role of host STING in immune protection is also reported in an E0771 tumor model after ionizing radiation (*12*). Without PolySTING treatment, faster MC38 tumor growth was observed in *Tmem173^−/−^* mice than wildtype mice, suggesting the presiding role of endogenous STING signaling in immune protection against immunogenic MC38 tumors. STING-mediated type I interferon (IFN-I), nuclear factor κB (NF-κB), and/or autophagy signaling are proposed to contribute to host defense against cancer or viral infection (*7, 13-15*). We evaluated the efficacy of PolySTING in mice deficient of the interferon-α/β receptor (*Ifnα/βR^-/-^*), and a STING mutant mouse (STING^S365A^) that blocks IRF3 binding and IFN-I production but not NF-κB activation or autophagy induction (*15*). Neither animal models exhibited antitumor efficacy upon PolySTING treatment over the wildtype mice (Fig. 3, E and F). Collectively, these data indicate that the host STING-IFN-I signaling is essential in PolySTING-mediated immunity against MC38 tumors.

**Fig. 3.**
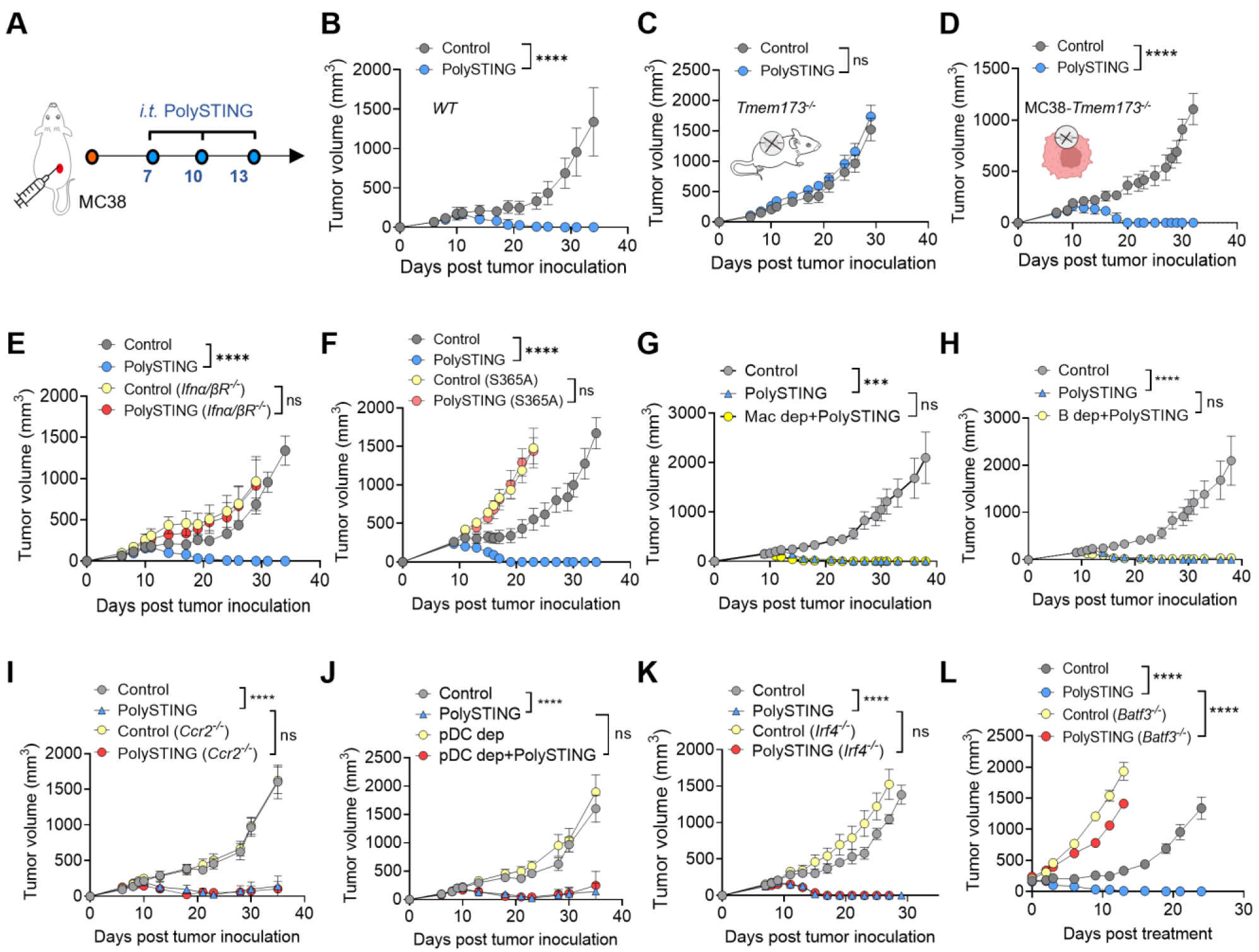
Identification of cDC1 for STING-mediated tumor rejection. (**A-D**) Tumor growth in wildtype, host STING knockout (*Tmem173*^-/-^) or cancer STING knockout (MC38-*Tmem173*^-/-^) C57BL/6 mice bearing MC38 tumors after PolySTING treatment. Tumor growth of (**E**) Ifnα/β receptor knockout (*Ifnα/βR^-/-^*) or (**F**) STING^S365A^ mutant mice bearing MC38 tumors after PolySTING treatment. Tumor growth in wildtype mice with antibody blockade of (**G**) macrophages (anti-CSF1R), (**H**) B cells (a mixture of anti-CD19/CD22/B220), and transgenic mice with depletion of different DC subtypes, including (**I**) MoDC (*Ccr2^-/-^*), (**J**) plasmacytoid DC (pDC, anti-CD317), (**K**) cDC2 (*Irf4^-/-^*), (**L**) cDC1 (*Batf3^-/-^*) mice, after PolySTING treatment. In all experiments, mice were intratumorally treated with PolySTING when the tumor volume reached around 200 mm^3^ in MC38 tumors. Data are represented as mean±sem, n≥6. Statical significance was calculated by two-way ANOVA, ns: no significant difference, ***: p<0.001, ****: p<0.0001.

### Identification of cDC1 for STING-mediated tumor rejection

To determine which immune cell subsets are important for PolySTING-mediated effect, we use antibody depletion or mice with deficiency in different subset of antigen presenting cells. MC38 tumor growth inhibition was evaluated by antibody blockade of B cells (a mixture of anti-CD19/CD22/B220), macrophages (anti-CSF1R), and in transgenic mice with depletion of different DC subtypes, including monocyte-derived DC (MoDC, *Ccr2^-/-^*) (*16*), plasmacytoid DC (pDC, anti-CD317) (*17*), cDC1 (*Batf3^-/-^*) (*18*), and cDC2 (*Irf4^-/-^*) (*19*). Depletions of B cells, macrophages, MoDCs, pDCs, and cDC2s did not exhibit a clear difference in antitumor efficacy over control mice after PolySTING treatment (Fig. 3, G to K). In contrast, mice with cDC1 deficiency (*Batf3^-/-^*) abolished the antitumor efficacy by PolySTING (Fig. 3L). Although free cGAMP did not display obvious cell tropism (fig. S1C), it still requires cDC1 to produce antitumor immunity, as the tumor growth inhibition is also abrogated in *Batf3^-/-^* mice (fig. S4A). Compared to free cGAMP, PolySTING has a stronger therapeutic effect in wildtype mice, likely due to the higher and more persistent STING signaling (Fig. 1B and fig. S1B) and increased uptake in myeloid cells (fig. S1, D to F).

To further determine whether the therapeutic contribution of cDC1 is dependent on endogenous STING activation or secondary to the IFN-I produced by neighboring cells, we constructed and investigated mice with conditional knockout of STING in cDC1 (*XCR1^cre^STING^fl/fl^*, fig. S4B) and monitored tumor growth after PolySTING treatment (Fig. 4A and fig. S4C). Tumor growth inhibition of MC38 tumors by PolySTING was abrogated in *XCR1^cre^STING^fl/fl^* mice, indicating STING signaling within cDC1s is required for the antitumor immunity.

**Fig. 4.**
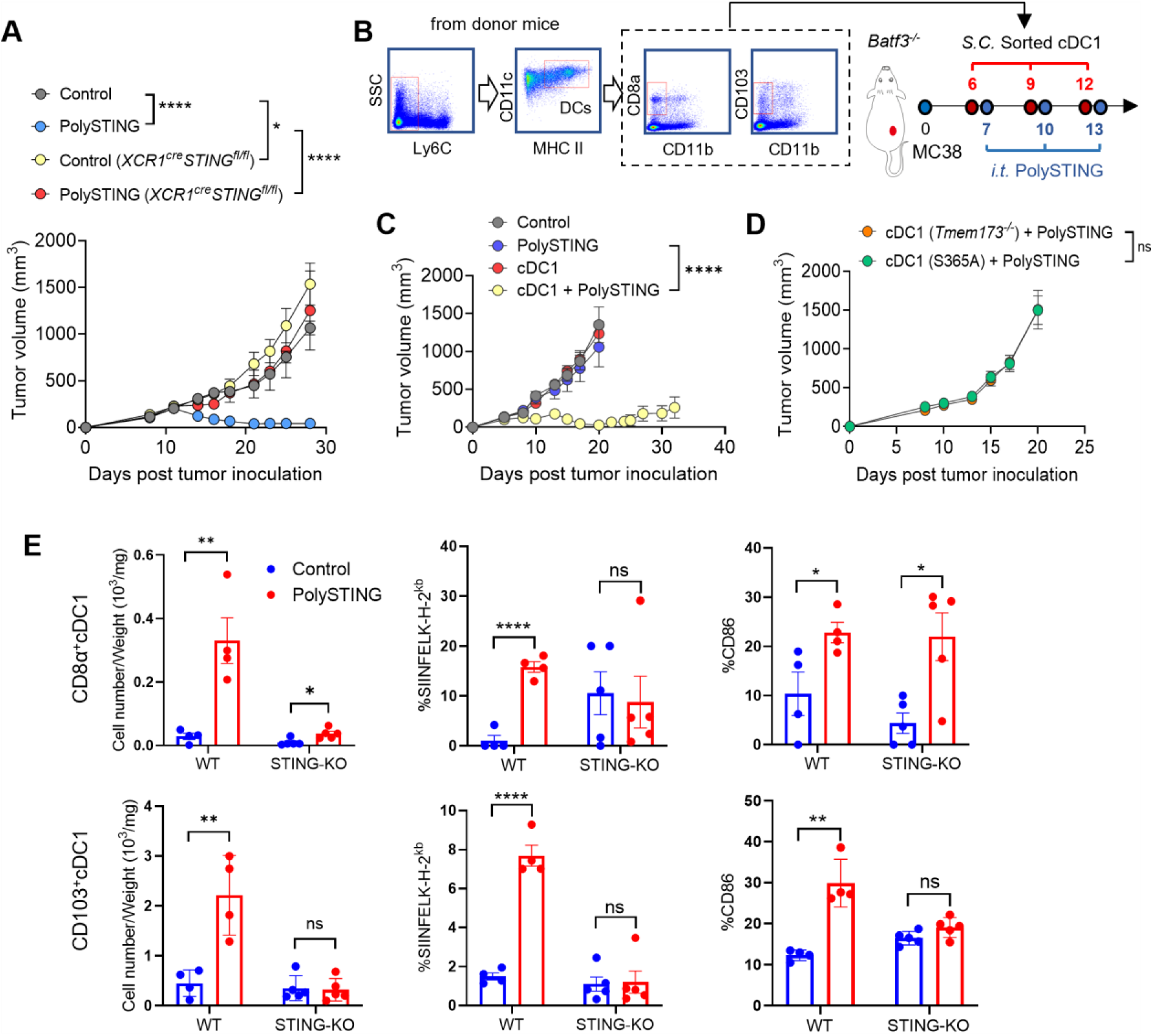
Direct STING signaling in cDC1 is essential for PolySTING-mediated antitumor immunity. (**A**) Tumor growth in conditional knockout of STING in cDC1s (*XCR1^cre^STING^fl/f^****^l^***) mice bearing MC38 tumors after PolySTING treatment. (**B-D**) MC38 tumor growth in *Batf3^-/-^* mice with or without PolySTING treatment after the mice were subcutaneously transferred with cDC1s from STING wildtype mice (C), STING knockout or STING^S365A^ mutant mice (D). Mice were intratumorally treated with PolySTING when the tumor volume reached around 200 mm^3^. cDC1s were sorted from BMDCs of donor mice. (**E**) Cell number per tumor weight, percentage of antigen (SIINFEKL) presentation and CD86 expression on CD8α^+^ cDC1s or CD103^+^ cDC1s with control or PolySTING treatment. Tumor tissues were harvested and analyzed by flow cytometry at 24 hours after treatments. All the data are represented as mean±sem, n=5. Statical significance was calculated by wo-way ANOVA in A-D, and two-tail t test in E. ns: no significant difference, *: p<0.05, **: p<0.01, ****: p<0.0001.

Next, we performed cell transfer studies in *Batf3^-/-^*mice to evaluate the sufficiency of cDC1 in STING-mediated antitumor efficacy. BMDCs were treated with GM-CSF, IL-4 and Flt3-L, and cDC1s were isolated and enriched through flow sorting (Fig. 4B). The combined CD8α^+^ and CD103^+^ cDC1s were subcutaneously injected adjacent to the MC38 tumors 12 hours prior to PolySTING treatment. Notably, adoptive transfer of cDC1s restored antitumor immunity and prolonged survival by PolySTING in *Batf3^-/-^* mice, whereas PolySTING treatment or cDC1 transfer alone showed negligible efficacy (Fig. 4C, and fig. S5, A and B). Transfer of cDC1s from STING knockout or STING^S365A^ mutant mice did not lead to immune protections against MC38 tumors in *Batf3^-/-^* mice (Fig. 4D). Lack of immune protection by cDC1 transfer from STING^S365A^ mutant mice supports previous findings that type I interferon is required by dendritic cells for immune rejection of tumors (*20, 21*).

To better define tumor specific responses, we utilized the MC38-OVA tumor model to investigate the priming of transferred cDC1s (wildtype versus STING knockout) in the *Batf3^-/-^*mice (Fig. 4E). In the MC38-OVA tumors, significant increase of both subsets of wildtype cDC1s (cell number was normalized to the weight of tumor) were observed with 10- and 4-fold higher CD8α^+^ and CD103^+^ cDC1s in PolySTING group over the glucose control, respectively. In contrast, STING deficient cDC1s show marginal difference between the two groups. Most notable increase is the OVA peptide (SIINFEKL) presentation on H-2K^b^ MHC-I complex for wildtype cDC1s, where PolySTING treatment elevated antigen presentation over 15- and 4-fold for CD8α^+^ and CD103^+^ cDC1s, respectively. In contrast, STING deficient cDC1s did not show any significant changes after PolySTING treatment in either subset of cDC1s. Co-stimulation of cDC1s (e.g., CD86) is also elevated in both subsets of cDC1s after PolySTING treatment. These results illustrate robust STING-dependent cDC1 infiltration and activation after PolySTING treatment.

### PolySTING boosts cDC1 activation and priming of CD8^+^ T cells *in vivo*

We further investigated STING-mediated potentiation of dendritic cells in antigen presentation, co-stimulation and CD8^+^ T cell priming after intratumoral injection in MC38-OVA tumors. MC38-OVA cells (1×10^6^ cells) were inoculated in C57BL/6 mice and grown to 150-200 mm^3^. On day 10 and 13, different doses of PolySTING (10, 50, 200 μg) were injected intratumorally, where tumors and tDLNs were harvested on day 14 for DC analysis and on day 16 for T cell analysis (fig. S6A). Notably, 50 μg of PolySTING induced the strongest cDC1 activation level and cytotoxic T cell response over the other two doses in both tumors and tDLNs (fig. S6, B to F), in accordance with prior therapeutic efficacy findings (fig. S1I).

We next performed a side-by-side comparison of PolySTING (50 μg) and Adu-S100 (50 μg) following a similar flow cytometry analysis (Fig. 5A). Results show significantly increased cDC1s in the MC38-OVA tumors from PolySTING group over either Adu-S100 (2.4-fold) or glucose control (7.1-fold) (Fig. 5B). Co-stimulation signals (e.g., CD80) were dramatically increased in both the CD8α^+^ and CD103^+^ cDC1s over Adu-S100 group (>15-fold, Fig. 5C). The percentage of dendritic cells with OVA peptide (SIINFEKL) presentation on H-2K^b^ MHC-I complex also significantly increased in the PolySTING group over the glucose control, but the difference with Adu-S100 group is insignificant (Fig. 5D). Dendritic cell activation by PolySTING resulted in significantly increased production of OVA-specific CD8^+^ T cells in both the tumor microenvironment and tumor-draining lymph nodes than the Adu-S100 and glucose groups (Fig. 5E). Tumor-resident CD8^+^ T cells are cytotoxic as demonstrated by the elevated interferon-γ and granzyme B expressions from PolySTING treatment (Fig. 5F). Combined with the previous data on Adu-S100-induced apoptosis in CD8^+^ T cells (Fig. 1E), these results highlight the importance of cell selectivity by PolySTING in mounting a productive antitumor immunity over known STING agonists such as Adu-S100.

**Fig. 5.**
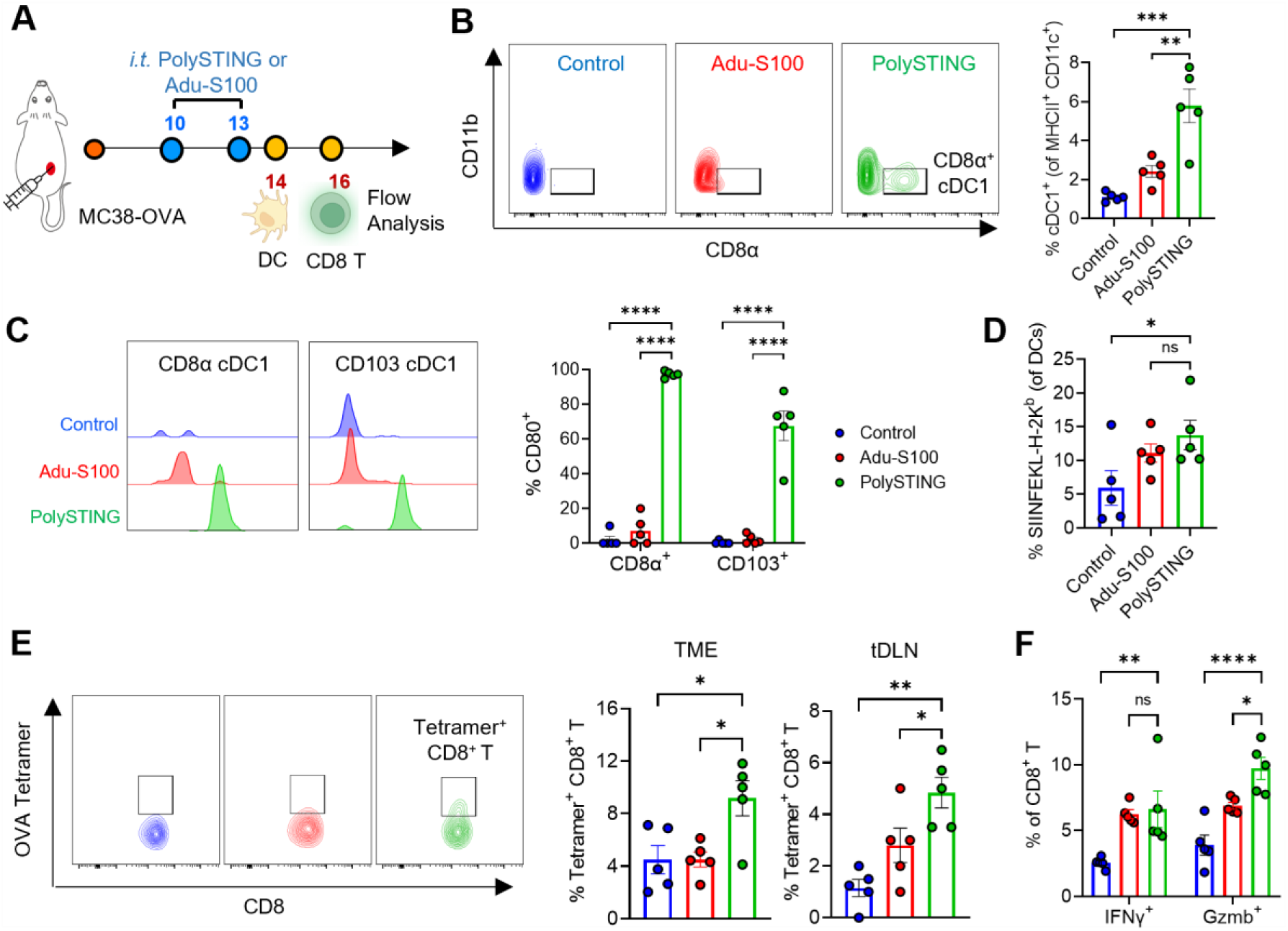
STING signaling boosts cDC1 activation and priming of CD8^+^ T cells. (**A**) Schematic overview of experimental design using MC38-OVA tumor bearing mice. PolySTING or Adu-S100 were intratumorally administered on day 10 and 13 when average tumor volumes reached 150 mm^3^. Dendritic cells and CD8^+^ T cells were harvested on day 14 and 16, respectively, for flow cytometry analysis. (**B**) Representative flow diagrams of CD8α^+^ cDC1 (gated on CD45^+^CD11c^+^MHCII^+^CD11b^-^) and quantification of percentage of cDC1 in total DC population. (**C**) Representative flow cytometric histograms of the costimulatory marker CD80 (gated on CD45^+^CD11c^+^MHCII^+^CD11b^-^CD8α^+^ or CD103^+^ cDC1) and quantification of CD80 percentage in CD8α^+^ and CD103^+^ cDC1s. (**D**) Percentage of antigen (SIINFEKL) presentation on H-2K^b^ complex of DCs in response to PolySTING or Adu-S100. (**E**) Representative flow cytometric plots of OVA-specific CD8^+^ T cells from the tumor and tumor-draining lymph nodes in response to PolySTING or Adu-S100 treatment. (**F**) Percentage of interferon-γ (IFNγ) and granzyme b (Gzmb) expressing CD8^+^ T cells in response to PolySTING or Adu-S100. All the data are represented as mean±sem, n=5. Statical significance was calculated by two-tailed t test, ns: no significant difference, *: p<0.05, **: p<0.01, ***: p<0.001, ****: p<0.0001.

Flow cytometry analysis shows CD103^+^ cDC1s from MC38 tumors also have significantly increased expressions of X-C motif chemokine receptor 1 (XCR1), C-type lectin domain containing 9A (CLEC9A) and C-C motif receptor 7 (CCR7) after STING activation (Fig. 6A). CCR7 is a cell surface chemokine receptor that drives DC homing to lymphoid organs (*22*). In the tumor draining lymph nodes, elevated CCR7 expressions were found on both subsets of cDC1s (Fig. 6B). *CCR7^-/-^* mice rejected cDC1 migration to lymph nodes and lost STING-mediated tumor growth inhibition (Fig. 6, C and D). These data demonstrate cDC1 trafficking to tumor-draining lymph nodes is important to prime the tumor-specific T cell immunity.

**Fig. 6.**
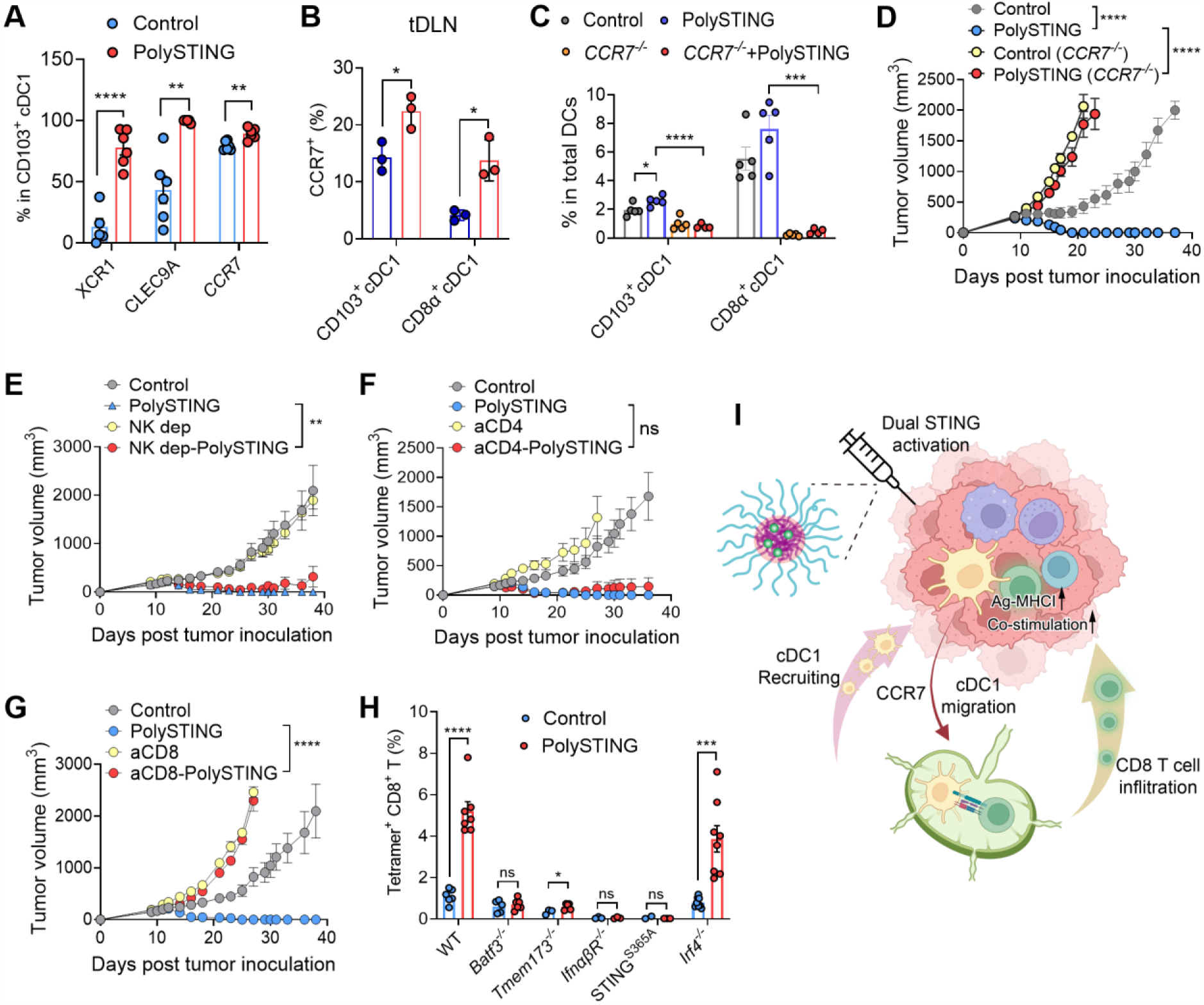
PolySTING-mediated migration of cDC1 and effect of T cells on antitumor efficacy. (**A**) Expression of XCR1, CLEC9A, and CCR7 on cDC1s in MC38 tumors 18 h after intratumoral PolySTING treatment. (**B**) Expression of CCR7 on cDC1s in tumor-draining lymph nodes (tDLN) after PolySTING treatment. (**C**) cDC1 percentage in total DCs in MC38 tumors and (**D**) tumor growth after PolySTING treatment in wildtype or CCR7 knockout (*CCR7^-/-^*) mice. MC38 tumor growth in C57BL/6 mice after depletion of (**E**) NK cells (anti-NK1.1), (**F**) CD4^+^ T cells (anti-CD4), (**G**) CD8^+^ T cells (anti-CD8a). (**H**) Tetramer^+^ CD8^+^ T cells from tDLN of wildtype and different genetic knockout mice with or without PolySTING treatment. (**I**) Schematic of STING-mediated cDC1 activation and priming of antigen-specific cytotoxic T cells. Mice receiving 5% glucose were used as controls. All the data are represented as mean±sem. n=6 in A,E-G, n=3 in B, n=5 in C,D, and n≥3 in H. Statical significance was calculated by two-tail t test in A-C,H, or two-way ANOVA in D-G. ns: no significant difference, *: p<0.05, **: p<0.01, ***: p<0.001, ****: p<0.0001.

Lastly, we depleted NK, CD4^+^ or CD8^+^ T cells and investigated their impact on PolySTING-mediated tumor clearance. Data revealed an indispensable role of CD8^+^ T cells but not NK or CD4^+^ T cells in tumor rejection (Fig. 6, E to G). Through an *in vivo* priming assay, MC38 tetramer^+^ CD8^+^ T cells yielded approximately 5-fold higher proliferation in the PolySTING treatment group over control in wildtype C57BL/6 mice (Fig. 6H). In contrast, only minimal tetramer^+^ CD8^+^ T populations were detected in *Batf3^-/-^, Tmem173^-/-^, Ifnα/βR^-/-^, or STING^S365A^* mice, whereas *Irf4^-/-^* (cDC2 knockout) mice maintained the production of antigen-specific CD8^+^ T cells. In summary, these data illustrate the important role of cDC1 activation by PolySTING in antigen presentation, co-stimulation, and migration to lymph nodes for the priming of CD8^+^ T cells against tumor cells (Fig. 6I).

### PolySTING-activated chemokine signature is associated with improved patient survival

We quantified the changes of immune profiles in the MC38 tumors in wildtype and cDC1 deficient (*Batf3^-/-^*) mice after STING stimulation. Tumors were harvested and homogenized 12 hours after a single injection of PolySTING, Adu-S100, or glucose control. CXCL9/10 and C-C motif chemokine ligand 4 and 5 (CCL4/5) are the top upregulated chemokines (>10-fold) in the wildtype mice after PolySTING treatment (Fig. 7A and fig. S7). In *Batf3^-/-^* mice, significantly reduced expression levels of CXCL9/10 and CCL4/5 were observed after PolySTING treatment, whereas such differences were insignificant in Adu-S100 groups (Fig. 7B). CXCL9/10 produced by cDC1s have been reported to drive adoptive T cell infiltration into tumors (*23*), and CCL4/5 upregulation in ovarian cancer patients has shown favorable response to checkpoint blockade therapy (*24*). Interestingly, Adu-S100 presents a different chemokine profile, wherein CXCL9 is downregulated while CCL20 and CXCL1 are upregulated (fig. S7A). These chemokines have been shown to increase the infiltration of Tregs and MDSCs, which tempers the inflamed tumor microenvironment (*25*).

**Fig. 7.**
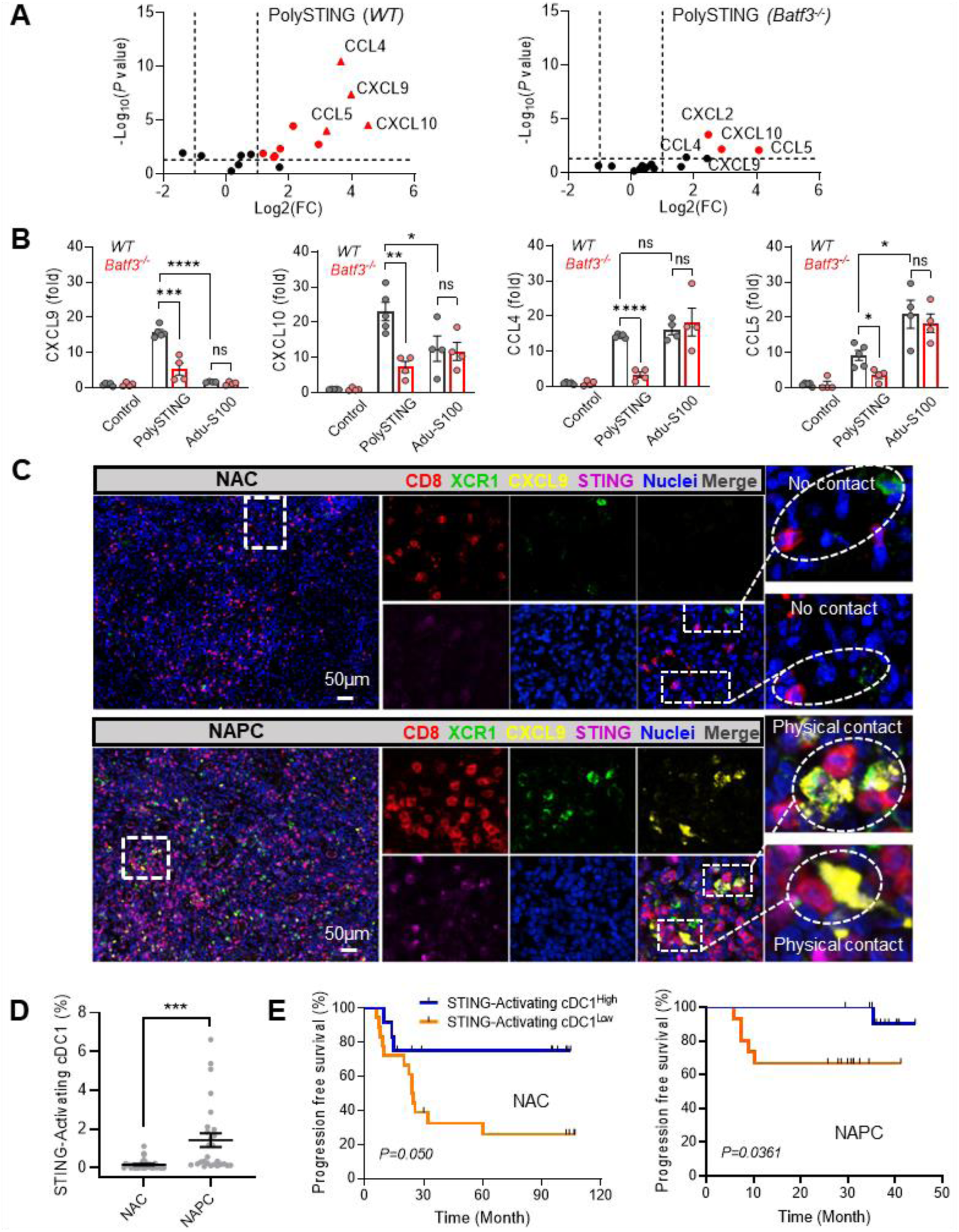
STING-activating cDC1 signature is associated with cancer patient survival. (**A**) Quantification of chemokine profile of MC38 tumors from wildtype or *Batf3^-/-^*mice after PolySTING treatment. (**B**) Comparison of expression levels of chemokines CCL4/5 and CXCL9/10 in wildtype versus *Batf3^-/-^* mice after indicated treatments. Data are represented as mean±sem, n≥4 for wildtype and *Batf3^-/-^* mice. (**C**) Representative images of mIHC staining of NSCLC tumors from either NAC or NAPC-treated patients. (**D**) Percentage of STING activating cDC1 was quantified and compared between NAC and NAPC-treated patients. (**E**) STING-activating cDC1 signature shows positive correlation with progression free survival in both NSCLC patients treated with either NAC or NAPC. Statical significance was calculated by two-tail t-test in B and D, and Logrank test was used to determine statistical significance for survival in E. *: p<0.05, **: p<0.01, ***: p<0.001, ****: p<0.0001.

Chemokines CCL5 and X-C motif chemokine ligand 1 (XCL1) from NK cells have been reported to induce cDC1 migration into tumors (*26*). In our study, RT-PCR analysis of MC38 tumors show significant increase in mRNA expressions of CCL4 and CCL5 but not XCL1 after PolySTING treatment (fig. S8A). An *in vitro* chemotaxis assay showed significant cDC1 enrichment in transwells containing CCL4 or CCL5, confirming their induction effect in cDC1 migration (fig. S8B). In mice, PolySTING expanded the infiltration of both subsets of cDC1 cells in the tumor microenvironment, while neutralization of CCL4/5 retarded STING-mediated cDC1 migration (fig. S8C) and tumor growth inhibition and survival (fig. S8, D and E).

We next examined The Cancer Genome Atlas (TCGA) database to evaluate PolySTING-induced chemokine signatures (CCL4/5, CXCL9/10) in cancer patients with skin cutaneous melanoma or sarcoma. In parallel, we also probed BATF3, CLEC9A, XCR1 and CLNK, a collective of biomarkers reported for cDC1s (*26*). Positive correlations were found between STING chemokine and cDC1 signatures in both sets of patient samples (Pearson correlation coefficient r>0.7, fig. S9, A and B). Furthermore, PolySTING-induced chemokine signatures (CCL4/5, CXCL9/10) are associated with improved survival outcomes in both cancer patient populations, whereas analysis by bulk STING expressions has no statistical significance (fig. S9, A and B.)

### STING-activated cDC1 signature is elevated in resected patient tumors after neoadjuvant checkpoint and chemotherapy

Based on the mechanistic findings from murine studies, we performed multiplex immunohistochemistry (mIHC) staining of formalin-fixed and paraffin-embedded tumor tissues from non-small cell lung cancer patients who have received either neoadjuvant chemotherapy (NAC, n=30) or neoadjuvant Pembrolizumab and chemotherapy (NAPC, n=28) (Table S1). We used XCR1^+^STING^+^CXCL9^+^ biomarkers to stain for STING-activating cDC1 cells, and further investigated their physical contact with CD8^+^ T cells in the tumor microenvironment (Fig. 7C and fig. S9C). Interestingly, patients who received NAPC therapy significantly elevated STING-activating cDC1 signatures over patients treated by chemotherapy alone (p<0.001, Fig. 7D), suggesting immune checkpoint therapy by Pembrolizumab is responsible for the cDC1 recruitment and activation in the tumor microenvironment in a subset of patients. The activated cDC1 further engages CD8^+^ T cells to augment the antitumor immunity. In accordance with mIHC analysis, we further divided the cancer patients into STING-activating cDC1^high^ (above medium level) and cDC1^low^ (below medium level) groups. STING-activating cDC1^high^ postulated a positive prediction of prolonged progression free survival and overall survival in both NAC and NAPC patients, whereas low abundance of STING-activating cDC1 signature led to poor survival outcome more pronouncedly in patients without immunotherapy (Fig. 7E and fig. S9D). These findings indicate that the STING-activating cDC1 signature (XCR1^+^STING^+^CXCL9^+^) may serve as a prognostic clinical biomarker for cancer patient response to therapies.

## DISCUSSION

Genomic instability of cancer cells places DNA sensing and cGAS-STING signaling as a key immune activation mechanism against cancer (*3*). However, immune system often displays dual host-protective and tumor-promoting responses that are antipodally activated to modulate immunity (*27*). This dichotomous relationship is illustrated by contradicting reports where STING signaling has opposite effect on antitumor response. For immune protection, DNA from dying cancer cells was shown to activate cGAS-STING pathway, leading to the induction of type I interferons that prime T cells and natural killer cells against tumor cells (*14, 28*). On the other hand, chromosome instability has also been reported to activate the cGAS-STING pathway with inflammatory responses that promote cancer metastasis (*29, 30*). Massagué *et al* show cGAMP transfer from tumor cells to astrocytes produces inflammatory cytokines which activate STAT1 and NF-kB pathways leading to tumor growth and chemoresistance (*30*). More recently, the same group reported an opposite effect where STING signaling in cancer cells suppresses metastatic outbreak in an indolent lung cancer model (*31*). STING is broadly expressed in majority of host cells including cancer cells, where multifaceted cellular functions likely exert complex functions in antitumor immunity. Therefore, understanding the cell context of STING activation is important to elucidate mechanisms that drive antitumor immunity.

Use of small molecule STING agonists (e.g., DMXAA, Adu-S100) for this purpose is challenging because they are readily taken up by T cells, and high magnitude of STING signaling has been shown to drive T cell deaths (*8, 32*) and ablate T cell memory response (*11*). To overcome this limitation, we utilized PolySTING, a nanoparticle STING agonist, to selectively target myeloid cells but not T cells, thereby preventing T cell deaths (Fig. 1, D and E). Prior studies have shown STING-activating nanoparticles are able to improve pharmacokinetics and cytosolic delivery of cyclic dinucleotides for STING activation (*33-35*). Irvine *et al* showed lipid nanodiscs increased dendritic cell infiltration in the MC38 tumors for T cell priming (*33*). Weichselbaum *et al* reported zinc cyclic di-AMP nanoparticles disrupted tumor endothelial cells and targeted macrophages for augmented antitumor immunity (*35*), whereas Wilson *et al* showed STING-activation nanoparticles led to vascular normalization with improved vascular integrity and T cell infiltration (*36*). In this study, our PolySTING design synergizes canonical STING activation by cGAMP with non-canonical activation by PSC7A, which prevents STING degradation and prolongs the expression of type I interferons (*10*). Using PolySTING and MC38 tumors as a model system, we systematically investigated the contributions of different immune cell subsets in tumor clearance. Data show depletion of cDC2, pDC, moDC, macrophages in the myeloid compartment or blocking of B and NK cells had minimal effect on tumor rejection (Fig. 3, G to K), whereas mice deficient of cDC1s (*Batf3^-/-^*) abrogated antitumor efficacy of PolySTING (Fig. 3L). This cDC1 requirement supports a recent study using cGAMP-loaded virus-like nanoparticles (*37*). Our data further demonstrate STING signaling within cDC1s is important because PolySTING efficacy is abolished in *XCR1^cre^STING^fl/fl^*mice (Fig. 4A), as well as transfer of cDC1s from STING knockout mice cannot rescue the abrogated PolySTING efficacy in *Batf3^-/-^* mice (Fig. 4C). Moreover, cDC1 recruitment in tumors harnesses an autocrine effect by CCL4/5 chemokines (*26*), which is highly induced by PolySTING in wildtype but not in *Batf3^-/-^* mice (Fig. 7, A and B). The collective chemokine signatures (CCL4/5, CXCL9/10) are further associated with better survival outcomes of melanoma and sarcoma patients over bulk STING expressions observed from the TCGA database (fig. S9, A and B). Additionally, high expressions of STING-activating cDC1s in tumor tissues from non-small cell lung cancer patients were observed in those treated with Pembrolizumab in combination with chemotherapy (NAPC) (Fig 7, C and D, and fig. S9C). The correlation between STING-activating cDC1^high^ and improved progression-free survival and overall survival were further validated (Fig. 7E and fig. S9D). For cancer patients who have low STING-activating cDC1 signatures even after immune checkpoint therapy (∼50% patients in Fig. 7D), PolySTING therapy may offer a synergistic treatment modality to boost cDC1 infiltration as demonstrated in the mouse tumor studies.

In mice with cDC1 deficiency, transfer of wildtype cDC1s but not STING deficient cDC1s restore antitumor immunity with PolySTING treatment (Fig. 4C). Recently, Murphy *et al* reported intratumoral injection of cDC1s but not cDC2s or GM-CSF-derived DCs led to rejection of immunogenic fibrosarcoma in mice lacking endogenous cDC1s (*38*). Using the MC38-OVA tumor model, our data further demonstrate PolySTING can boost transferred cDC1s in antigen presentation and co-stimulation (Fig. 4E). Recent data show early priming of CD4^+^ T cells against tumor antigens also requires cDC1s, and in turn, cognate CD4^+^ T cell interactions and CD40 signaling are required to license cDC1 for the activation of CD8^+^ T cells (*38*). PolySTING-cDC1-mediated antitumor immunity bypasses the co-licensing requirement of CD4^+^ T cells as blocking of CD4^+^ T cells had little effect on tumor clearance (Fig. 6F). Further studies are warranted to elucidate the molecular and cellular mechanisms of sufficiency in direct STING activation of cDC1s for CD8^+^ T cell priming.

In summary, using a nanoparticle STING agonist, we discovered the importance of STING licensing of conventional type I dendritic cells (cDC1s) over other immune cell subsets in mounting a robust antitumor immunity. A STING-activating cDC1 signature (XCR1^+^STING^+^CXCL9^+^) may serve as a prognostic clinical biomarker for patient response to cancer immunotherapy. Our findings indicate direct targeting of STING pathways in the cDC1s offers an effective strategy to restore protective immunity against solid tumors.

## MATERIALS AND METHODS

### Study design

The primary objective of this study was to use a “shock-and-lock” STING-activating nanoparticle (PolySTING) as a tool to uncover new biological insights on cell context dependence of STING-mediated antitumor immunity. Conventional small molecule agonists (2’3’-cGAMP or Adu-S100) were used as a treatment of reference. The tumor growth and animal survival of different tumor-bearing mice were monitored. Mice of specific gene knockout or conditional knockout were genotyped, while mice of specific immune cell blockade were confirmed via flow cytometry before each experiment. Through loss (*XCR1^cre^STING^fl/fl^*)- and gain (cDC1 adoptive transfer)-of-function studies, we demonstrated direct STING signaling in cDC1 is both necessary and sufficient for the rejection of multiple established and metastatic murine tumors. The immune orchestration of cDC1 and CD8^+^ T cells in tumors/tDLNs by different treatments was carefully characterized and the chemokine/cytokine profiles of tumors were quantified in wildtype and cDC1-deficient mice. We further collected tumor tissues from non-small cell lung cancer patients who have received NAC or NAPC treatment, and performed mIHC staining to evaluate the clinical value of STING-activating cDC1 signature. Sample size was determined via a priori power analysis based on means and SEMs estimated from pilot experiments or the literature. The number of samples combined and the number of independent experiments are included in the figure legends.

### Patient participants and ethical approvals

A total of 58 non-small cell lung cancer (NSCLC) patients who received two cycles of neoadjuvant chemotherapy (NAC, 30 cases) or neoadjuvant pembrolizumab chemotherapy (NAPC, 28 cases) prior to surgical resection were included in this study. The squamous cell carcinoma patients received neoadjuvant paclitaxel 175 mg/m^2^ plus carboplatin (area under curve 5; 5 mg/mL per min) and the adenocarcinoma patients received pemetrexed 500 mg/m^2^ plus carboplatin, with or without intravenous pembrolizumab 200 mg on day 1, 21 days each cycle, for two cycles before surgical resection, and then followed by two cycles after surgical resection. Clinicopathological parameters included age, gender, smoking index, clinical tumor node metastasis (TNM) stage, pathological type. Progression-free survival (PFS) was defined as the time from the date of surgery to the date of recurrence or last follow-up. Overall survival (OS) was defined as time from the date of surgery to the date of any caused death or last follow-up. The last follow-up was conducted on October 10, 2022, with a median follow-up time of 97 and 35 months for NAC and NAPC patients, respectively. The study was approved by the Ethics Committee of Tianjin Cancer Institute & Hospital (protocol number: bc2020060). All patients enrolled in the study provided written informed consent.

### Cell lines and culture

*Tmem173^-/-^* MC38 cells were generated by knocking out *Tmem173* genes by CRISPR/Cas9 technology following a published procedure (*39*). LLC2 cells were kindly provided by Zhijian ‘James’ Chen, UT Southwestern Medical Center. TC-1 cells were kindly provided by T. C. Wu, Johns Hopkins University. 4T1-luciferase cells were kindly provided by S. Huang, Massey Cancer Center, Virginia Commonwealth University. MC38, MC38-OVA, and B16F10 cells were purchased from ATCC (Manassas, VA). All the cells were cultured in complete DMEM medium (Gibco-BRL Life Technologies, Canada) supplemented with 10% fetal bovine serum (FBS) (HyClone, USA), 100 U/ml penicillin, and 100 mg/ml streptomycin (Gibco-BRL Life Technologies, Canada). THP1-ISG cells from ATCC were cultured in RPMI medium supplemented with 10% FBS and 0.05 mM β-mercaptoethanol (β-ME). All cells were grown at 37 °C in a humid atmosphere with 5% CO_2_. Bone marrow cells were filtered through a 70 μm strainer and treated with Red Blood Cell Lysis Buffer (Sigma). Single-cell suspensions of bone marrow cells were cultured in RPMI-1640 medium containing 10% FBS, supplemented with 20 ng/ml GM-CSF and 10 ng/ml IL-4 for BMDC, or 20 ng/ml M-CSF for BMDM (Peprotech). The medium was half replenished every 2 days. Cell lines were routinely monitored for mycoplasma contamination. All cells used in this study tested negative for mycoplasma contamination.

### Mice

*XCR1^cre^* and *STING^fl/fl^* mice were purchased from the Jackson Laboratory. The *STING^fl/fl^* mice were first crossed with *XCR1^cre^* mice to generate F1 mice. Then the F1 mice were inbred to generate the cre^+^ mice with homozygous *STING^fl/fl^*. The insertion of cre and flox elements were confirmed by PCR per Jackson Laboratory’s protocol. *Tmem173^-/-^* mice, *IfnαβR^-/-^*mice, *Ccr2^-/-^* mice, *Batf3^-/-^* mice, *Irf4^-/-^*mice, *Ccr7^-/-^* mice were purchased from the Jackson Laboratory. C57BL/6 wild-type (*WT*) mice, BALB/c mice were purchased from Charles River Laboratories. All the mice were maintained under specific pathogen-free conditions and housed in a barrier facility under a 12 h–12 h light–dark cycle and maintained on standard chow (2916, Teklad Global). The temperature range for the housing room is 68–79 °F (average is around 72 °F) and the humidity range is 30–50% (average is around 50%). The experimental groups included randomly chosen female littermates of ages around 8 weeks and the same strain. All experiments were performed in accordance with the procedures approved by the AAALAC-accredited Institutional Animal Care and Use Committee at UT Southwestern Medical Center and were compliant with all relevant ethical guidelines.

### Preparation of PolySTING nanoparticles

PEO-*b*-P(MAC-SC7A·HCl) (PSC7A) copolymer was synthesized following a previously published procedure (*40*). cGAMP-PSC7A nanoparticles (PolySTING) were prepared by mixing 25 μL 2′3′- cGAMP (2 mg/mL) in 1 mL PSC7A polymer solution (1 mg/mL) containing 5% D-glucose at pH 5, and then adjusted to pH 7 using NaOH (1 M) to induce micelle assembly with cGAMP loading. After micelle formation, the nanoparticles were characterized using dynamic light scattering (Zeta Sizer, Malvern, He-Ne laser, λ = 633 nm) to measure the hydrodynamic size, size distribution and zeta potential. Transmission electron microscopy was used to analyze particle morphology. The cGAMP loading efficiency (>90%) was quantified via high-performance liquid chromatography (1260 Infinity II, Agilent).

### Microscopic imaging of cell uptake of PolySTING

BMDCs and BMDMs were grown in a four-well glass chamber and treated with fluorescently labelled PolySTING (cGAMP-FITC encapsuled in Cy5-labeled PSC7A) for the indicated time. Cells were fixed in 4% paraformaldehyde, then stained for CD11c (Alexa Fluor 594 anti-mouse CD11c antibody, clone: N418, BioLegend), or CD11b (Alexa Fluor 594 anti-mouse CD11b antibody, clone: M1/70, BioLegend) using an immunofluorescence kit (Cell Signaling Technologies). Samples were stained with DAPI (Thermo Fisher Scientific) and imaged using the built-in software (ZEN 2.6) of the Zeiss 700 confocal laser scanning microscope with a 63× oil-immersion objective. ImageJ v.1.52d was used to quantify colocalization using the Pearson’s correlation coefficient. Data are representative of at least 20 cells.

### Tumor growth and treatments

For subcutaneous tumor xenografts, C57BL/6 mice were inoculated with 1×10^6^ MC38 colon carcinoma cells, 1×10^6^ LLC2 lung cancer cells, 1.5×10^5^ TC-1 tumor cells, or 1.5×10^5^ B16F10 melanoma cells on the right flank on day 0. The metastatic breast cancer model in Balb/c mice was constructed by mammary fat pad injection of 1×10^6^ 4T1-luciferase cancer cells. MC38 tumor-bearing C57BL/6 mice were treated by intratumoral injection of PolySTING (50 μL per mice) Adu-S100, or 5% glucose control on day 11, 14 and 17. B16F10 tumor-bearing mice were treated on day 8, 11 and 14. TC-1 tumor-bearing mice were treated on day 7, 10 and 13. LLC2 tumor bearing mice were treated on day 6, 9 and 12. 4T1 tumor-bearing mice were treated with PolySTING on day 11, 14 and 17. Meanwhile, anti-PD-1 (200 μg) was intraperitoneally injected one day before PolySTING and was given 3 times. Mice were monitored for tumor growth. Tumor volumes were measured with a caliper by the length (L), width (W) and calculated as V = L×W×W/2.

For distal tumor model, C57BL/6 mice were subcutaneously injected with 1×10^6^ MC38 cells or 2×10^5^ B16F10 cells on the primary site, followed by subcutaneous injection of 1×10^6^ MC38 cells on day 4 or 2×10^5^ B16F10 cells on day 6 on the contralateral sites. Mice were intratumorally injected with PolySTING on day 10, 13 and 16. Anti-PD-1 was given one day before PolySTING. Mice were monitored for tumor growth on both primary and distal sites. For MC38 rechallenge study, the tumors of C57BL/6 wildtype mice were removed by surgical resection on day 30 and then mice were injected with 1×10^6^ MC38 cells on distal site on day 45. Meanwhile, C57BL/6 naïve mice were injected with MC38 cells on day 45. Cured mice from PolySTING treatment were rechallenged with the same amount of MC38 cells on primary site and distal site. Mice were monitored for tumor growth.

### In *vivo* immune cell depletion experiments

For the NK cell depletion assay, mice were i.p. injected with anti-NK1.1 antibody (500 μg, clone PK136, BioXcell) on day 3 after tumor inoculation and then were maintained with 250 μg of anti-NK1.1 antibody every 3 days to deplete NK cells. For CD8^+^ or CD4^+^ T cell depletion, mice were treated with anti-CD8a antibody (200 μg, clone YTS169.4, BioXcell) or anti-CD4 antibody (200 μg, clone GK1.5, BioXcell) by i.p. injection on day 3 after tumor inoculation and then were maintained with same dosage of antibodies every 3 days. For B cell depletion, mice were i.p. treated with a cocktail of antibodies (*41*), including CD19 (clone 1D3, BioXcell), B220 (clone RA3.3A1/6.1, BioXcell), and CD22 (clone CY34.1; BioXcell) weekly, each at 150 mg per mouse. After 48 h, the mice were injected with a secondary antibody (mouse anti-rat Kappa light chain, clone MAR 18.5, BioXcell) at 150 mg. In addition, mice were injected at 250 mg weekly for 4 weeks. For macrophage depletion, mice were treated with 500 μg of anti-CD115 antibody (clone AFS98, BioXcell) every 3 days. Plasmacytoid dendritic cells (pDC) depletion in C57BL/6 mice was achieved by i.p. injection of mice with 200 μL of anti-PDCA1 antibody (clone 927, BioXcell) at 2 mg/mL.

### Adoptive transfer of dendritic cells

BMDCs were generated by culturing bone marrow cells flushed from the femurs of C57BL/6 naïve mice, STING knockout mice, or STING^S365A^ mutant mice. The RPMI 1640 medium supplemented with 10% FBS, Pen/Strep, sodium pyruvate, 20 ng/mL GM-CSF, 10 ng/mL IL-4 and 200 ng/mL Flit-3L was prepared for culturing BMDCs. Non-adherent and loosely adherent immature DCs were collected on day 6 and induction efficacy of BMDCs was detected by determining the expression of CD11c (routinely 70%–80% CD11c^+^). CD103^+^ cDC1 and CD8α^+^ cDC1 were sorted by flow cytometry (*42*) and then mixed as total cDC1. Then 5×10^5^ cDC1s were subcutaneously injected into MC38 tumor-bearing *Batf3^-/-^* mice by the tumor one day before PolySTING treatment. Mice were monitored for tumor growth.

### Flow cytometry

Tumor tissue or tumor-draining lymph nodes (tDLN) were harvested from mice with indicated treatment and minced using surgical scissors. Tissues were then digested using 160 μg/mL collagenase Ⅳ (Sigma– Aldrich) and 20 μg/mL DNase Ⅰ (Sigma–Aldrich) in RPMI 1640 media at 37 ℃ for 30 minutes and then washed and strained through a 70 μm filter (BD Falcon). LIVE/DEAD™ Fixable Aqua Dead Cell Stain Kit (Invitrogen™, L34966), PerCP anti-mouse CD45 (clone 30-F11, BD), Brilliant Violet 650 anti-mouse Ly-6C Antibody (clone HK1.4, BioLegend), Brilliant Violet 421 anti-Mouse CD11b (clone M1/70, BD), Brilliant Violet 786 anti-mouse CD11c (clone HL3, BD), PE/Cyanine7 anti-mouse I-A/I-E (clone M5/114.15.2, BioLegend), BV605 anti-mouse CD103 (clone M290, BD), Alexa Fluor 700 anti-mouse CD8a (clone 53-6.7, BioLegend), APC anti-mouse CD86 (clone GL-1, BioLegend), PE/Cyanine 7 anti-mouse CD80 (clone 16-10A1, BioLegend), PE anti-mouse XCR1 (clone ZET, BioLegend), PE anti-mouse CLEC9A (clone 7H11, BioLegend), PE/Cyanine 7 anti-mouse CCR7 (clone 4B12, BioLegend), Brilliant Violet 786 anti-mouse CD3e (clone 145-2C11, BD), Alexa Fluor 700 anti-mouse CD8 (clone 53-6.7, BioLegend), Brilliant Violet 605 anti-mouse CD4 (clone RM4-5, BioLegend), APC anti-mouse IFN-γ (clone XMG1.2, BioLegend), FITC anti-mouse Granzyme B (clone GB11, BioLegend), PE anti-mouse H-2K^b^ bound to SIINFEKL (clone 25-D1.16, BioLegend) APC anti-mouse T-Select H-2Kb MuLV p15E Tetramer-KSPWFTTL (code TB-M507-2, MBL) were used for flow cytometry analyses. Cell suspensions were incubated with Fc Block TruStain FcX Clone 93 (BioLegend) in PBS containing 0.5% bovine serum albumin and 2 mM EDTA before staining with fluorochrome labeled antibodies, followed by staining with live/dead dyes and selective antibodies of cell surface markers. Intracellular markers, including IFN-γ and granzyme B, were stained after cell fixation and permeabilization with True-Nuclear^TM^ transcription factor buffer set per protocol (BioLegend). Data were collected on BD LSR Fortessa or Beckman CytoFLEX flow cytometer and analyzed by FlowJo (Tree Star Inc., Ashland, OR) software.

### Tetramer^+^ CD8^+^ T cell assay

C57BL/6 wildtype mice as well as different types of genetic knockout mice (*Batf3^-/-^, Tmem173^-/-^, IfnαβR^-/-^, STING^S365A^, Irf4^-/-^*) were subcutaneously inoculated with 1ⅹ10^6^ MC38 cancer cells on the right flank on day 0. Mice were then treated with PolySTING when the tumor volume reached 150-200 mm^3^. Tumor draining lymph nodes from these mice were made into single cell suspensions the second day after two injections of PolySTING. The ratio of tetramer^+^ CD8^+^ T cells with or without PolySTING treatment from mice were analyzed by flow cytometry.

### Multiplex cytokine and chemokine analysis

Fresh MC38 tumor tissue from wildtype or *Batf3^-/-^* C57BL/6 mice with PolySTING, Adu-S100, or 5% glucose control treatment were analyzed for levels of cytokine/chemokine expressions by choosing mouse cytokine/chemokine 44-plex discovery assay array (MD44). Tumor tissue homogenate preparation was performed according to the manufacturer’s instructions (www.evetechnologies.com).

### Tissue isolation and quantitative PCR analysis

Fresh MC38 tumor tissues were homogenized by using Precellys Evolution machine (Bertin). Total RNAs were extracted from tumor tissues using the RNeasy mini kit (Qiagen). RNA quantity and quality was then confirmed using the NanoDrop (DeNovix DS-11) system. Genomic DNA was removed and cDNA was synthesized using an iScript gDNA clear cDNA synthesis kit (Bio-Rad). Bio-Rad SsoAdvanced universal SYBR green supermix and CFX connect real-time system were used for PCR analysis. Results were corrected by GAPDH and plotted in GraphPad Prism version 9.0. The DNA primers used were as follows: mouse Ccl4: AGCCAGCTGTGGTATTCCTG, AGGGGCAGGAAATCTGAACG; mouse Ccl5: CTCACCATATGGCTCGGACA, CTTCTCTGGGTTGGCACACA; mouse Xcl1: CTTTCCTGGGAGTCTGCTGC, CAGCCGCTGGGTTTGTAAGT.

### Dendritic cell chemotaxis assay

Chemotaxis of cDC1s was analyzed by using transwell migration assays. BMDCs (1×10^6^ cells) were dispersed in RPMI medium with 1% BSA and seeded into 5 μm pore size transwell inserts (Corning) of a 24-well tissue culture plate containing 500 ml RPMI/BSA medium with CCL4 (R&D) and CCL5 (R&D) at a final concentration of 150 ng/mL. After incubation for 3 h, migrated CD103^+^ cDC1 or CD8α^+^ cDC1 cells in the lower compartment were collected and quantified by flow cytometry. Migration was calculated as fold change between CCL4/5-treated group over the control group.

### Bioinformatic analysis of cancer patient data

RNA-seq by expectation-maximization (RSEM)-normalized expression datasets from The Cancer Genome Atlas (TCGA) were used. Hierarchical clustering of expression data was plotted as heatmaps using the ‘gplots’ package (version 3.0.1), where red indicates higher and blue indicates lower expressions relative to the mean expression per gene. For generation of gene expression signatures, normalized expression values were log2-transformed and ranked by the mean expression value of signature genes. The following gene signatures were used: STING expression (*Tmem173*), PolySTING-activated chemokines (CCL4, CCL5, CXCL9, CXCL10), cDC1 (CLEC9A, XCR1, CLNK, BATF3). Overall survival analyses were performed for the top and bottom quartile expression ranked values for selected genes or the ranked sum expression of gene signature and plotted for Kaplan-Meier curves using GraphPad Prism version 9.0.

### Multiplex immunohistochemistry and multispectral analysis

Formalin-fixed and paraffin-embedded (FFPE) tumor tissues from patients post-NAC or -NAPC were collected for multiplex immunohistochemistry (mIHC) staining. mIHC staining was performed using a PerkinElmer Polaris 7-color Technology Kit according to the manufacturer’s instructions. Spectra Systems was used to scan and photograph multispectral images at 20× magnification with the same exposure times with 10 randomly fields in each stained slides selected. The multispectral images were split and segmented using InForm image analysis software. Individual cells were identified with a nuclear segmentation algorithm using DAPI staining, together with a cellular mask around each nucleus to quantitate surface marker expression for each cell.

### Statistical analysis

Statistical analysis was performed using Microsoft Excel and Graphpad Prism Version 9.0. Values reported in figures are expressed as the standard error of the mean, unless otherwise indicated. For normally distributed datasets, we used 2-tailed Student’s t test and two-way ANOVA followed by Bonferroni’s multiple comparison test. For survival analysis, p values were calculated using the Log Rank test. p values > 0.05 were considered not significant (ns), p values < 0.05 were considered significant. * p value < 0.05, ** p value < 0.01, *** p value < 0.001, **** p value < 0.0001.

### Data availability

The main data supporting the results in this study are available within the paper and the Supplementary Information. All data generated in this study, including source data and the data used to generate the figures, are available from the corresponding authors upon request.

## Supplementary Materials

**This PDF file includes:**

Figs. S1 to S9

Table S1

**Other Supplementary Material for this manuscript includes the following:**

MDAR Reproducibility Checklist

## Acknowledgements

We thank T.C. Wu (Johns Hopkins) for providing the TC-1 cells, Z.J. Chen (UT Southwestern) for LLC2 cells, and S. Huang and Q. Chang (VCU Health) for 4T1-luciferease cells. We are grateful for suggestions and technical supports from Gao lab members, especially Z. T. Bennett, Q. Feng, J. Chen, W. Li and J. Wilhelm. This work was supported by grants from the National Institutes of Health (U54 CA244719, R01CA216839 to J.G.), National Natural Science Foundation of China (U20A20375 to J.W.), Cancer Prevention and Research Institute of Texas (RP220150 to J.G.) and Mendelson-Young Endowment in Cancer Therapeutics.

## Author contributions

J.W. and S.L. planned the study under the guidance of J.G., performed majority of the experiments, and analyzed the data. M.W. established the *XCR1^cre^STING^fl/fl^* mice with the help of Z.S. and performed the DC-T cell priming experiments in the MC38-OVA tumor model. X.W. carried out PSC7A synthesis and PolySTING formulation. S.C. performed bioinformatic analyses and some RT-PCR measurements. X.R. contributed to the clinical sample collection and mIHC methodology. Y.X.F. and N.Y. assisted with multiple gene-knockout and immune-cell-dependent animal studies. S.L., J.W., M.W., G.H., B.D.S., Y.X.F. and J.G. prepared and revised the manuscript.

## Data and materials availability

All materials used or generated in this study are available to researchers following appropriate standard material transfer agreements. All data are available in the main text or the supplementary materials.

