## Supplementary Materials for "STING Licensing of Type I Dendritic Cells Potentiates Antitumor Immunity"

**This PDF file includes:**

Figs. S1 to S9
Table S1

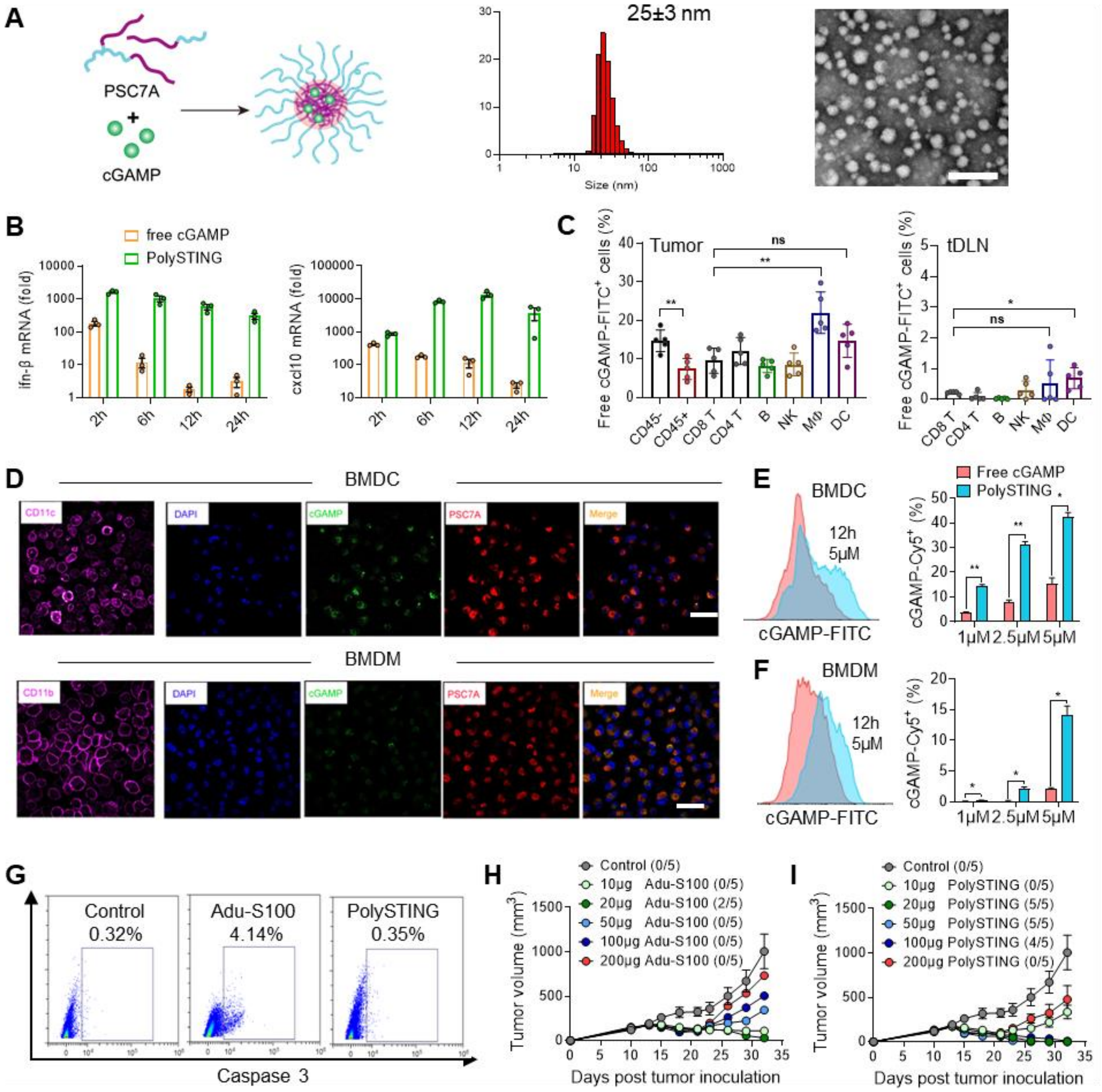


**Fig. S1. Physical characterization, cell tropism, and dose-response of PolySTING nanoparticle and free STING agonist.** (**A**) Synthetic scheme, size distribution and transmission electron micrograph of PolySTING nanoparticles. (**B**) qPCR measurement of ifn-β/cxcl10 mRNA expressions in THP1 cells after free cGAMP or PolySTING treatment at indicated time points. (**C**) Percentage of $FITC^+$ cells amongst cell populations in the MC38 tumors and tDLNs following intratumoral administration of FITC-labeled cGAMP. (**D**) Confocal microscopy and (**E,F**) flow cytometry of FITC-cGAMP-loaded Cy5-labeled PSC7A nanoparticle uptake by BMDCs and BMDMs after 12 h incubation at different concentrations. (**G**) Flow cytometric plots of T cell apoptosis (gated on $CD45^+CD3^+CD8^+$Caspase $3^+$) induced by Adu-S100 compared to PolySTING treatment and control. (**H,I**) MC38 tumor growth curves after intratumoral injection of Adu-S100 or PolySTING at indicated doses. The number of tumor free mice was indicated in brackets. Data are represented as mean±sem, n≥3. Statical significance was calculated by two-tail t-test. *: p<0.05, **: p<0.01. Scale bar, 100 nm in A, and 50 μm in D.

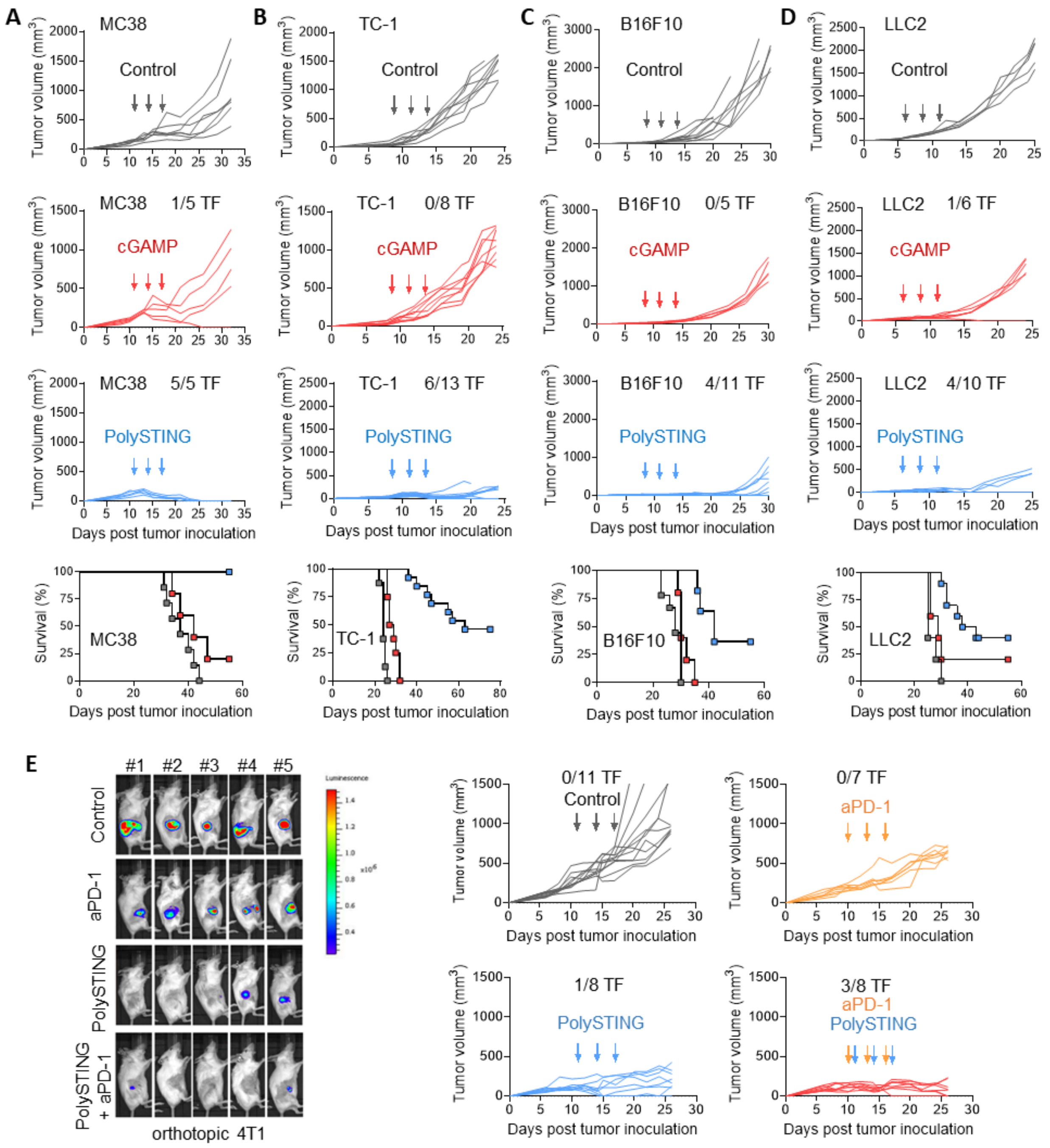


**Fig. S2. PolySTING induces antitumor immunity against established and metastatic tumor models.** (**A-D**) Individual tumor growth and overall survival after intratumoral administration of PolySTING, free cGAMP, or glucose control in MC38, TC-1, B16F10, or LLC2 tumors in C57BL/6 mice. TF: tumor free. The survival data are represented as mean±sem. Statical significance was calculated by or Mantel–Cox tests, ****: p<0.0001. (**E**) Individual orthotopic 4T1 tumor growth image of luciferase signal on week 3 and primary tumor growth curve after different treatments. The representative images of luciferase signal on week 3 are shown here, while lung metastasis data are shown in Fig. 2E.

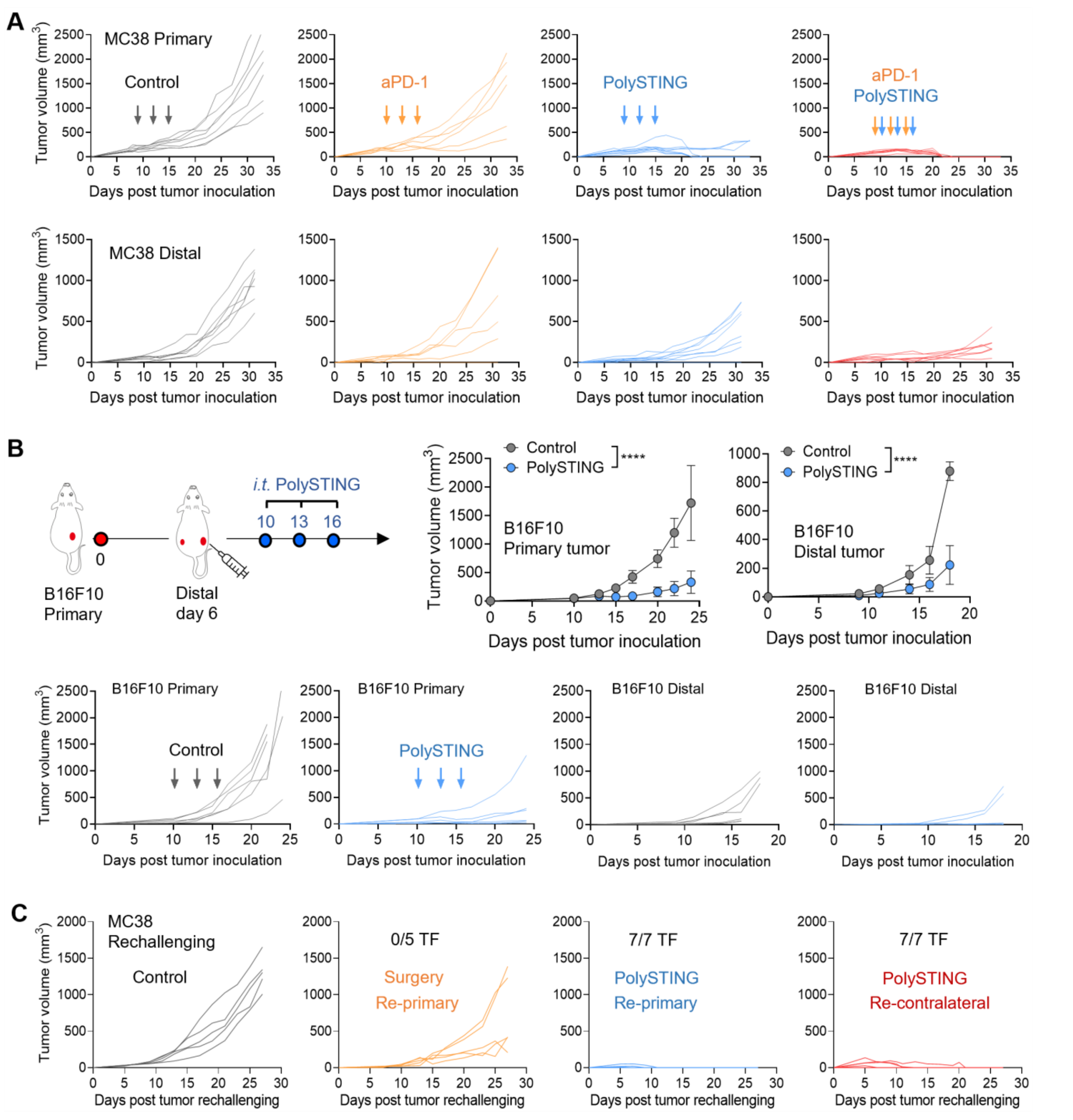


**Fig. S3. PolySTING induces antitumor immunity against distal tumors and tumor rechallenging.** (**A**) Individual tumor growth of injected primary and untreated contralateral tumor in C57BL/6 mice bearing MC38 upon indicated treatment. (**B**) Injected primary and untreated contralateral tumor growth in C57BL/6 mice bearing B16F10 tumors upon indicated treatment. Data are represented as mean±sem, n=6. Statical significance was calculated by two-way ANOVA, ****: $p<0.0001$. (**C**) Individual tumor growth after MC38 tumor rechallenging.

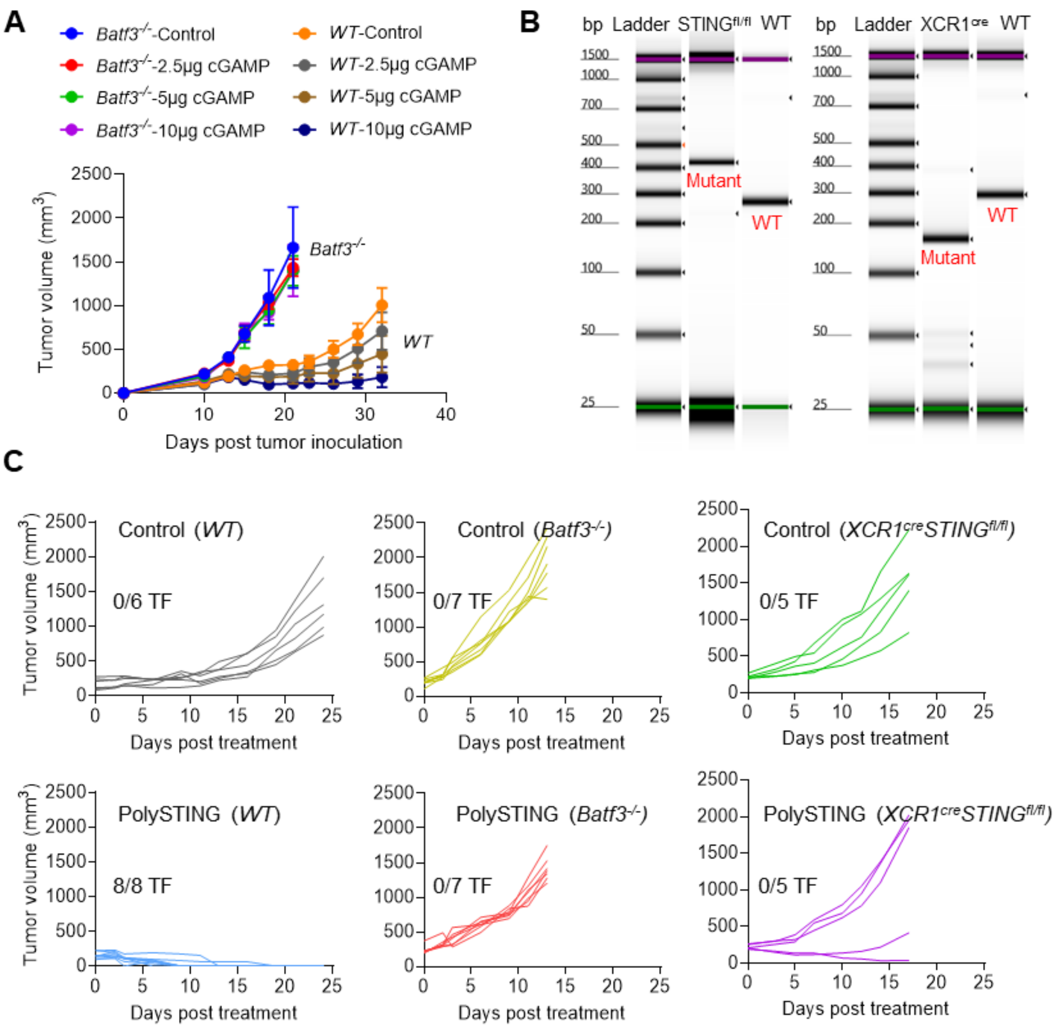


**Fig. S4. Identification of cDC1 for STING-mediated tumor rejection.** (**A**) MC38 tumor growth in wildtype or cDC1-deficient (*Batf3*$^{-/-}$) mice after different doses of cGAMP treatment. Data are represented as mean±sem, n=5. Statical significance was calculated by two-way ANOVA, *: p<0.05, ****: p<0.0001. (**B**) Genotype analysis of condition knockout mice (*XCR1*$^{Cre}$ *STING*$^{fl/fl}$). PCR product of homozygous floxed STING is at ~450bp, and the size of XCR1 cre product is at 287bp for wildtype and ~160bp for mutant. Homozygous flox and heterozygous cre genotype show both products in one condition knockout mouse. C57BL/6 wildtype mice are used as negative control. (**C**) Individual MC38 tumor growth after intratumoral administration of PolySTING or glucose control in C57BL/6 mice with indicated gene knockout. TF: tumor free.

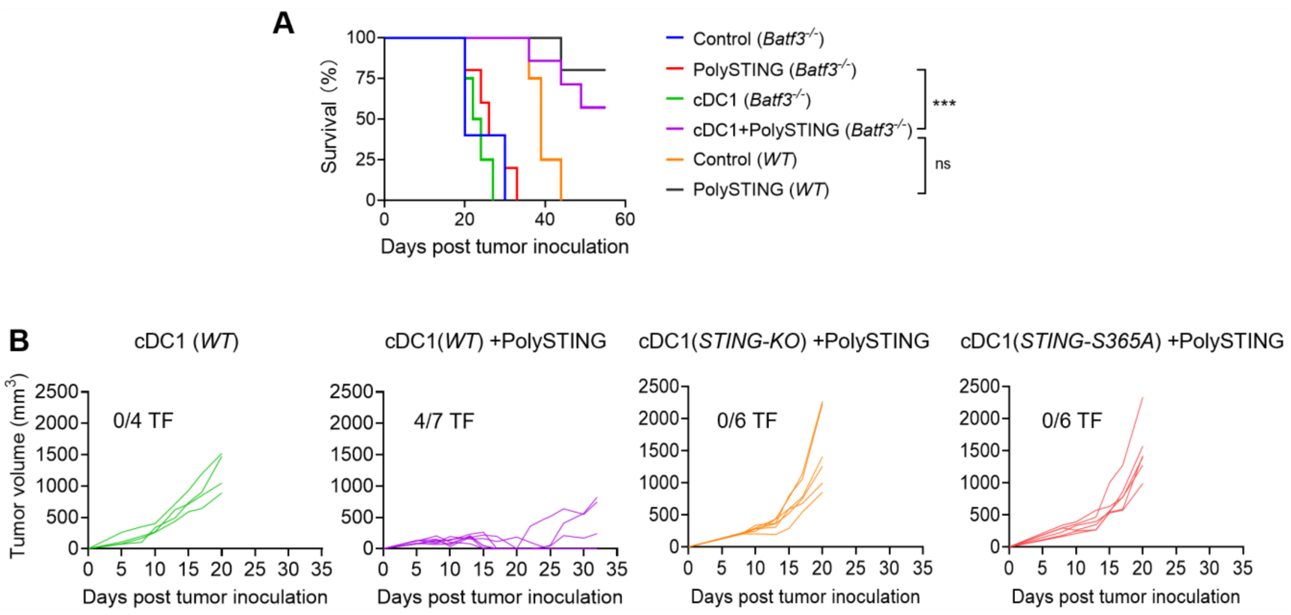


**Fig. S5. PolySTING primes wildtype but not STING-deficient transferred cDC1s in *Batf3*$^{-/-}$ mice.** (**A**) Kaplan–Meier survival curves upon indicated treatments in MC38 tumor-bearing wildtype or *Batf3*$^{-/-}$ C57BL/6 mice. TF: tumor free. The survival data are represented as mean±sem, n≥5. Statical significance was calculated by or Mantel–Cox tests, ****: p<0.0001. (**B**) Individual MC38 tumor growth upon intratumoral PolySTING treatments and subcutaneously transferred cDC1s from STING wildtype, knockout, S365A mutant mice.

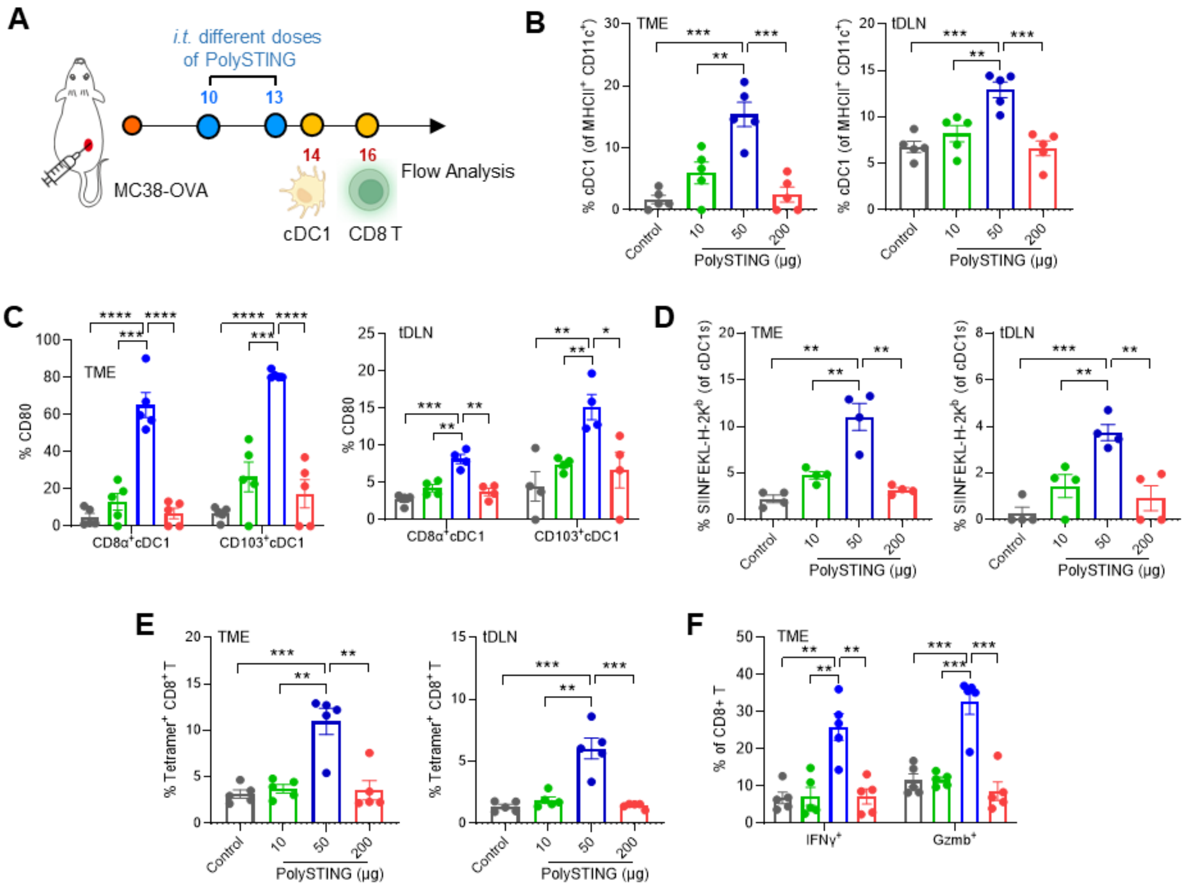


**Fig. S6. Dose-response of cDC1 activation and priming of CD8$^+$ T cells by PolySTING.** (**A**) Schematic overview of experimental design using MC38-OVA tumor bearing mice. Different doses of PolySTING (10, 50, 200 μg per mouse) or 5% glucose control were intratumorally administered on day 10 and 13 when average tumor volumes reached 150 mm$^3$. cDC1 and CD8$^+$ T cells were harvested on day 14 and 16, respectively, for flow cytometry analysis. (**B**) Quantification of percentage of cDC1 in total DC population. (**C**) Quantification of CD80 percentage in CD8α$^+$ and CD103$^+$ cDC1s. (**D**) Percentage of antigen (SIINFEKL) presentation on H-2K$^b$ complex of cDC1s. (**E**) Percentage of OVA-specific CD8$^+$ T cells from the tumor and tumor-draining lymph nodes. (**F**) Percentage of interferon-γ (IFNγ) and granzyme b (Gzmb) expressing CD8$^+$ T cells. All the data are represented as mean±sem, n=5. Statical significance was calculated by two-tail t-test. *: $p<0.05$, **: $p<0.01$, ***: $p<0.001$, ****: $p<0.0001$.

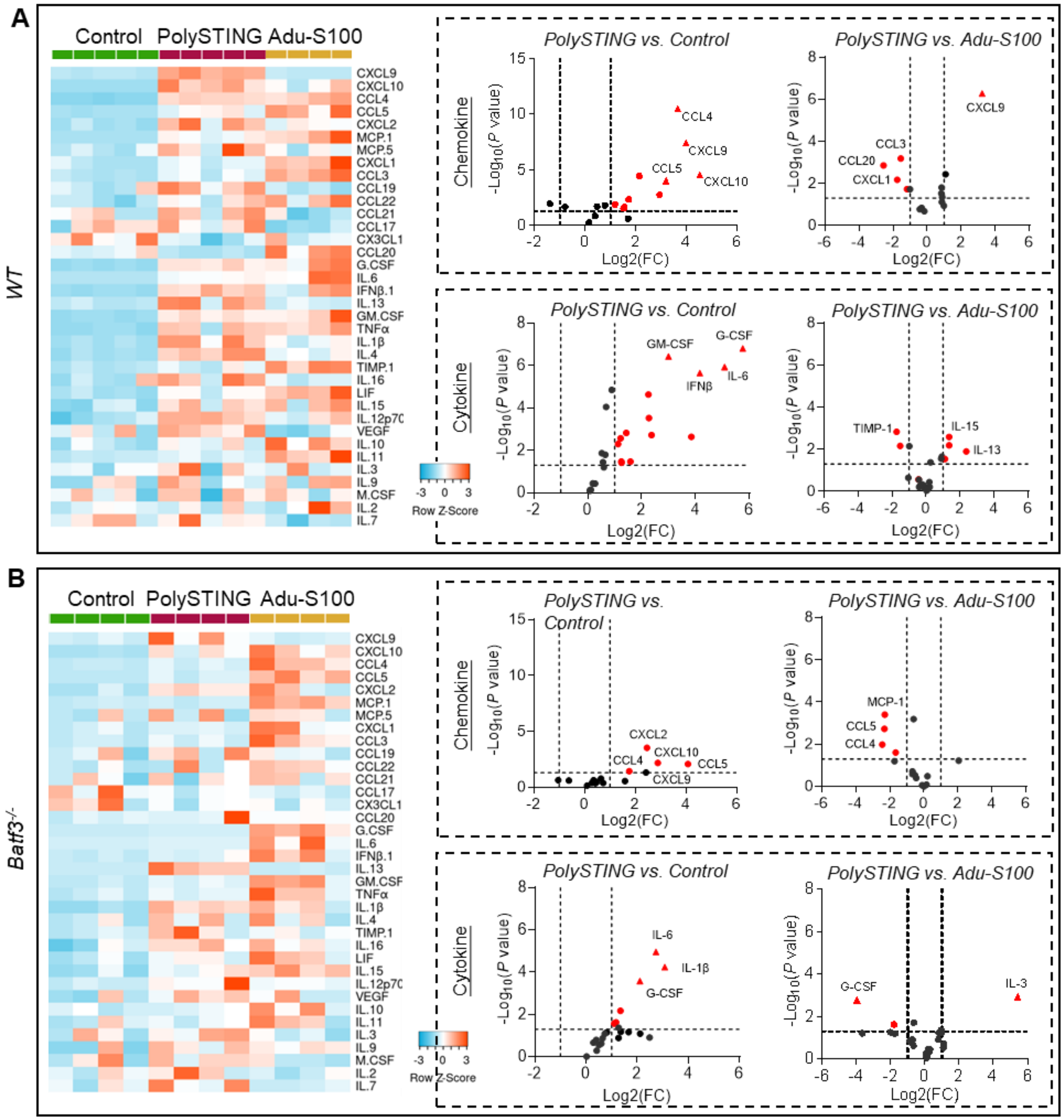


**Fig. S7. Changes of immune profiles in MC38 tumors after PolySTING or Adu-S100 treatment in wildtype and cDC1 deficient mice.** (**A,B**) Quantification of chemo and cytokine profiles of MC38 tumors in (A) wildtype or (B) *Batf3*$^{-/-}$ mice after 50 µg PolySTING, Adu-S100, or 5% glucose treatment. n≥4.

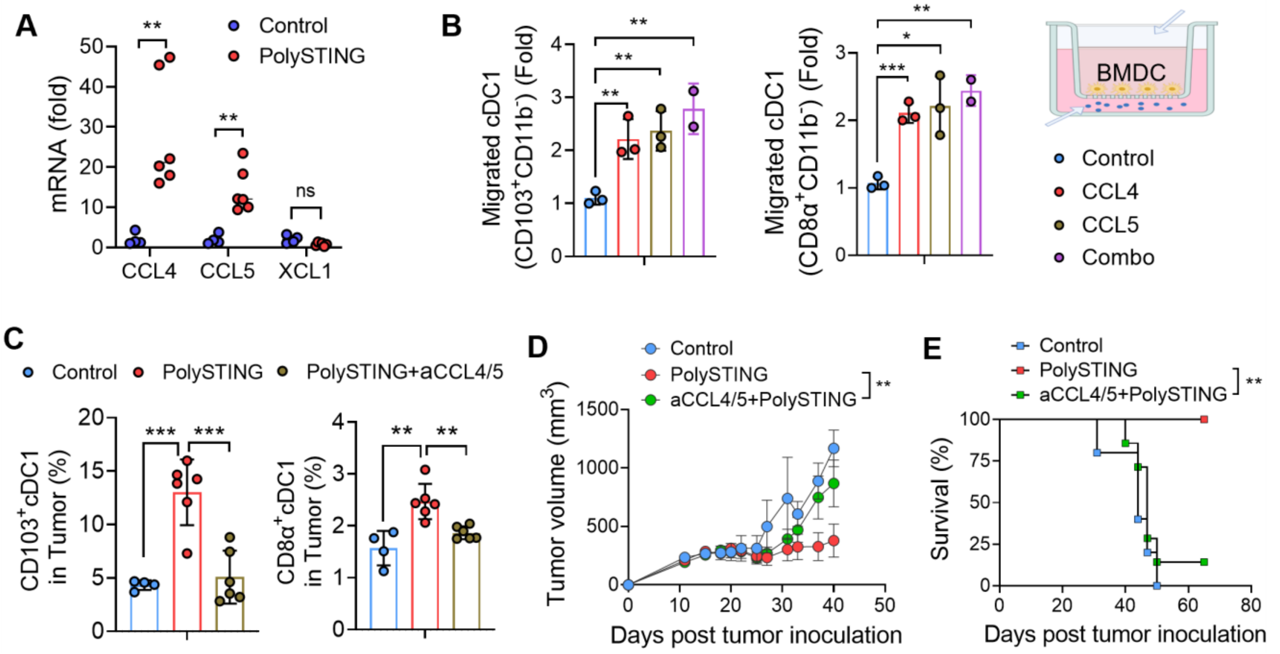


**Fig. S8. PolySTING mediated cDC1 recruitment and tumor growth inhibition in a CCL4/5-dependent manner.** (**A**) mRNA expression of CCL4, CCL5, and XCL1 in MC38 tumor tissues after PolySTING or 5% glucose treatment. (**B**) Chemotaxis analysis of both subsets of cDC1s towards CCL4/5 (150 ng/ml) in transwell migration assays. (**C-E**) Blockade of CCL4/5 reduced cDC1 infiltration in MC38 tumors (C) and abrogated STING-mediated tumor growth inhibition (D,E). CCL4 and CCL5 neutralizing antibodies (50 µg each) were intravenously injected on day 3 and maintained every 3 days in MC38 tumor-bearing mice. All the data are represented as mean±sem, n≥5 in A,C-E, and n=3 in B. Statical significance was calculated by two-tail t-test in A-C, two-way ANOVA in D, or Mantel–Cox tests in E. ns: no significant difference, *: $p<0.05$, **: $p<0.01$, ***: $p<0.001$.

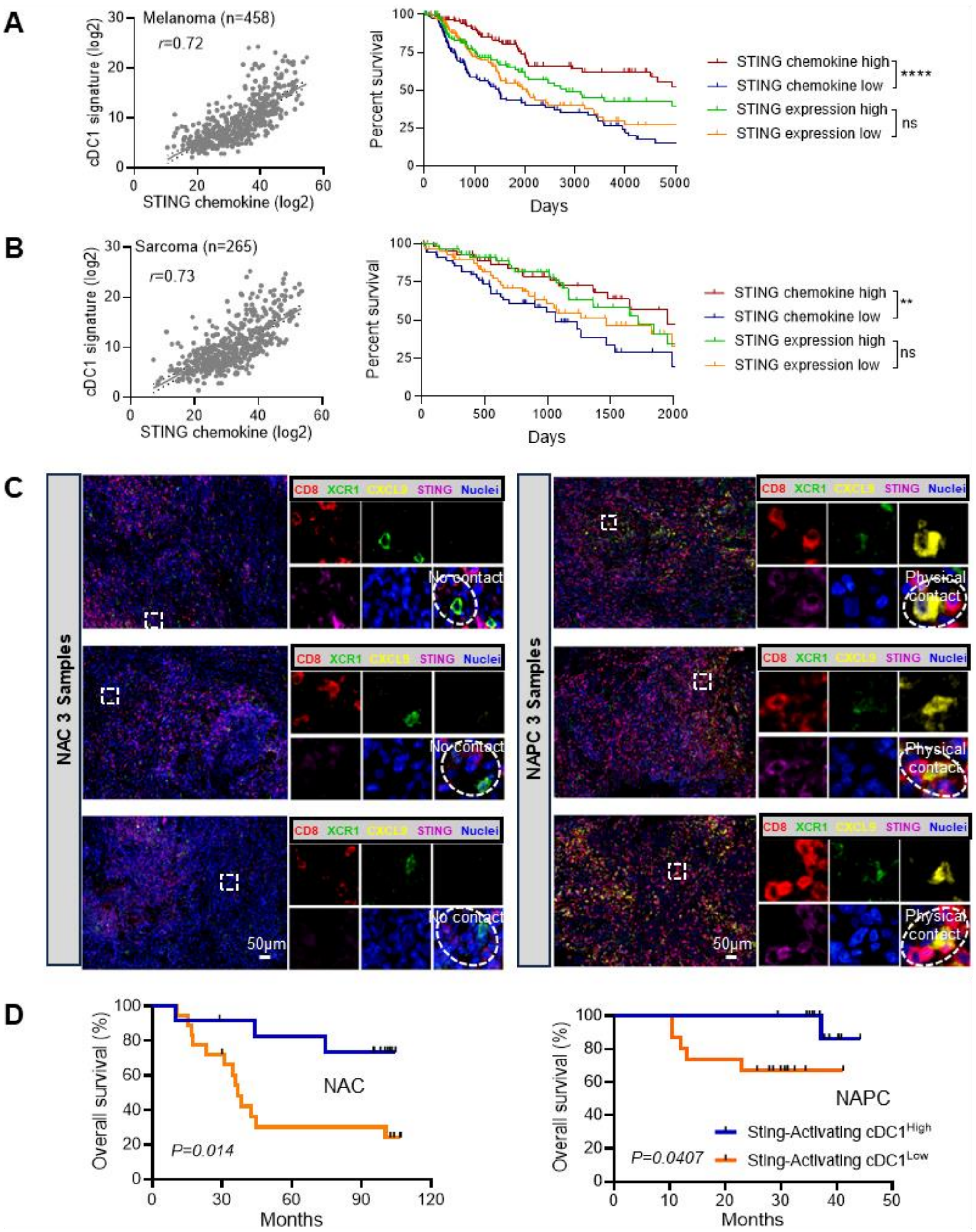


**Fig. S9. STING-activating cDC1signature displays positive correlations with patient survival.** STING activated chemokine expression levels (CCL4/5, CXCL9/10) positively correlates with cDC1 biomarkers (BATF3, CLEC9A, XCR1, CLNK) and offers higher prognostic value than STING expressions for survival in (**A**) melanoma or (**B**) sarcoma patients. A total of 458 melanoma patients or 265 sarcoma patients from TCGA datasets were analyzed, and top and bottom quartiles were compared. (**C**) Three additional representative images of multiplex immunohistochemistry staining of NSCLC tumors from either NAC or NAPC patients, respectively. (**D**) STING-activating cDC1signature show positive correlation with overall survival in both NSCLC patients treated with either NAC (30 patients) or NAPC (28 patients). The Logrank test was used to determine statistical significance for overall survival, ns: no significant difference, **: p<0.01, ****: p<0.0001.

**Table S1.** Patient and pathological characteristics of non-small cell lung cancer patients treated with neoadjuvant chemotherapy (NAC) or neoadjuvant Pembrolizumab and chemotherapy (NAPC).

| **Characteristics** | | **NAC (N=30)** | **NAPC (N=28)** | ***P* value** |
|---|---|---|---|---|
| Age (year) | | | | 0.17 |
| | ≤60 | 13 | 7 | |
| | >60 | 17 | 21 | |
| Gender | | | | 0.9 |
| | Male | 26 | 24 | |
| | Female | 4 | 4 | |
| Smoking index | | | | 1.0 |
| | ≤400 | 15 | 14 | |
| | >400 | 15 | 14 | |
| Pathology | | | | 0.69 |
| | Squamous | 17 | 19 | |
| | Adenocarcinoma | 11 | 7 | |
| | Large cell | 2 | 1 | |
| | Sarcomatoid | 0 | 1 | |
| Stage | | | | 0.6 |
| | I | 1 | 0 | |
| | II | 6 | 5 | |
| | III | 23 | 23 | |

*P values (>0.05) indicate no significant biases in the selection of patients based on the clinical and pathological characteristics
